# Retrosplenial PV and SST interneurons shape egocentric spatial precision and stability

**DOI:** 10.64898/2026.05.10.724096

**Authors:** Daehyun Oh, Jiyeon Yang, Jaehan Shin, Jeehyun Kwag

## Abstract

Accurate navigation requires egocentric representations of environmental geometry to be continuously updated by self-motion while remaining stable over time. The retrosplenial cortex (RSC) is central to this process, yet how local inhibitory circuits support this balance remains unclear. We show that parvalbumin (PV) and somatostatin (SST) interneurons regulate distinct components of egocentric spatial coding in RSC. PV interneurons are strongly modulated by self-motion and exhibit bearing-aligned synchrony that precedes SST activation, linking movement to egocentric coding precision. In contrast, SST interneurons display weak self-motion modulation but robust boundary-anchored activity with globally coherent dynamics that stabilize representations over time. Optogenetic silencing revealed dissociable effects: PV perturbation degraded egocentric coding precision while SST perturbation disrupted global population organization. Behaviorally, PV silencing impaired initial egocentric orientation while SST silencing preserved initial orientation but impaired its sustained update. These findings identify separable inhibitory mechanisms balancing rapid updating with representational stability during navigation.

## Introduction

Accurate navigation requires the brain to continuously update internal representations of space as the body moves through the environment. This process depends on integrating self-motion signals—including angular head velocity, linear speed, and movement trajectory—with information about environmental geometry to preserve a coherent sense of orientation ^1, 2^. While hippocampal–entorhinal circuits encode allocentric representations anchored to external landmarks ^3, 4^, successful navigation also relies on egocentric representations that track spatial relationships relative to the moving body ^5, 6^.

The retrosplenial cortex (RSC) is critical for this function. Disruption of RSC impairs heading orientation, route planning, and navigation across species^7–9^. Positioned between hippocampal–thalamic spatial circuits and parietal–motor association cortices, RSC has been proposed to mediate transformations between allocentric and egocentric reference frames required for navigation ^6, 10–12^. Consistent with this role, excitatory neurons in RSC encode egocentric boundaries, environmental vertices, and heading-relative geometry, with population activity mapping environmental structure in body-centered coordinates ^13–19^.

Egocentric spatial coding presents a fundamental computational challenge. Because the reference frame itself moves with the body, spatial representations must be updated with every rotation or translation. At the same time, unchecked updating would lead to cumulative error and representational drift. Theoretical and experimental work suggests that spatial circuits resolve this tension by combining rapid self-motion–driven updating with stabilizing mechanisms that constrain population activity over longer timescales ^6, 20^. How such a balance between flexibility and stability is implemented at the level of cortical microcircuits remains unclear.

Inhibitory interneurons are well positioned to regulate this balance. Parvalbumin (PV) interneurons provide fast, perisomatic inhibition that sharpens spike timing, synchronizes neuronal populations, and supports temporally precise network dynamics ^21–24^. In hippocampal and entorhinal circuits, PV inhibition contributes to the temporal organization of place and grid cell firing and supports accurate path integration ^25–27^. Somatostatin (SST) interneurons, in contrast, provide dendrite-targeting inhibition that regulates synaptic integration and modulates population activity over longer timescales, contributing to representational stability and memory persistence ^28–30^.

Despite extensive work on inhibitory specialization in hippocampal–entorhinal circuits, the role of inhibitory interneurons in retrosplenial spatial coding remains poorly understood. Interneurons in RSC have been implicated in contextual memory and network synchronization ^31–33^, yet whether inhibitory circuits in RSC participate directly in self-motion processing or egocentric spatial coding has not been established. How inhibitory microcircuits organize, update, or stabilize egocentric representations in RSC therefore remains unknown.

Here, we address this gap using cell-type–specific calcium imaging, optogenetic perturbations, and population-level analyses in freely navigating mice. We show that PV and SST interneurons in RSC are functionally segregated by spatial reference frame and temporal dynamics. PV interneurons strongly couple to self-motion signals, encode egocentric boundary orientation, and exhibit fast, bearing-aligned synchrony that precedes SST activation. In contrast, SST interneurons show weak self-motion modulation but robust boundary-anchored activity and globally coherent dynamics that stabilize spatial representations over time. At the population level, PV inhibition sharpens the precision of egocentric excitatory states, whereas SST inhibition preserves their global organization and long-timescale stability. These findings identify distinct inhibitory circuit mechanisms controlling the precision and stability of egocentric spatial coding during navigation in RSC.

## Results

### PV and SST interneurons encode environmental geometry using distinct spatial reference frames

To determine whether RSC PV and SST interneurons encode environmental geometry, we injected AAV-DIO-GCaMP6s in RSC of PV-Cre and SST-Cre mice and performed one-photon calcium imaging of GCaMP6s-expressing PV and SST interneurons using miniaturized microscopes during free exploration (Fig. 1a). GCaMP6s expression in PV and SST interneurons was confirmed and Ca^2+^ dynamics were recorded, from which spikes were inferred by deconvolution (Fig. 1b).

**Fig. 1:**
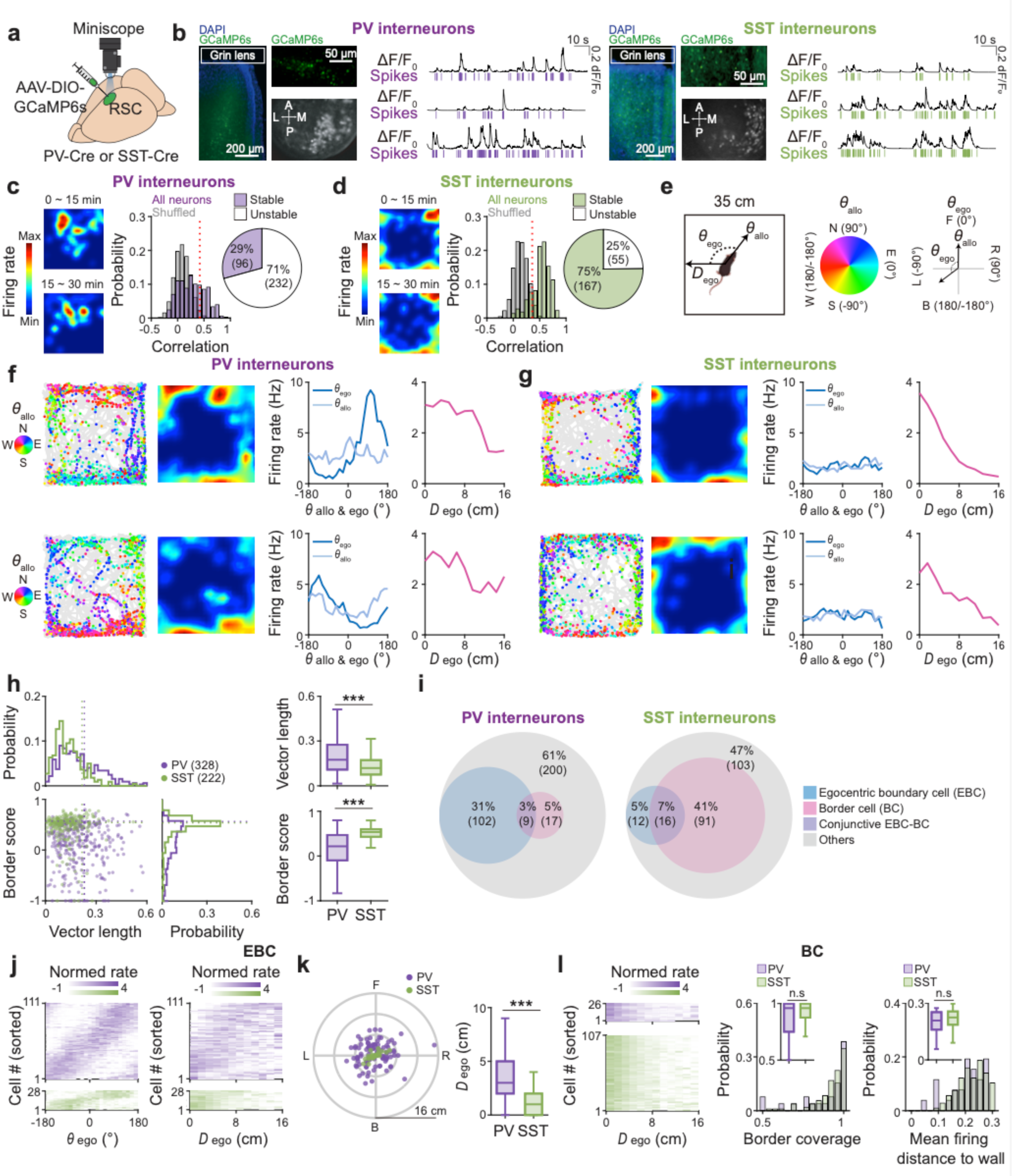
Distinct spatial reference frames encoded by PV and SST interneurons in the retrosplenial cortex. **a**, Schematic of the experimental setup for AAV-DIO-GCaMP6s injection, GRIN lens implantation, and miniaturized microscope-based Ca² imaging in the retrosplenial cortex (RSC) of PV-Cre and SST-Cre mice. **b**, Representative field of view (FOV) of Ca² imaging (left) and representative raw Ca² signals with corresponding deconvolved spikes (right). **c-d**, Representative spatial firing rate maps during the first and last 15-min of the 30-min free exploration in a square open chamber (left), probability distributions of spatial stability correlations (right) for PV (**c**, purple) and SST interneurons (**d**, green). Vertical red dotted line: 99^th^ percentile of randomly shuffled distribution. **e**, Illustration of allocentric head direction (θ_allo_), egocentric bearing (θ_ego_), and egocentric distance (*D*_ego_) to the closest boundary relative to the animal’s heading direction (left). The θ_allo_ of each spike is shown using a color-coded polar plot (middle) and the corresponding θ_ego_ rfor each spike is shown elative to θ_allo_ (right). **f-g**, Representative spike-trajectory plots (left; mouse trajectory: gray line, spike locations: colored dots) and corresponding firing rate maps, θ_ego_ tuning curves (middle, dark blue) and allocentric head direction θ_allo_ tuning curves (middle, light blue), and egocentric distance (*D*_ego_) tuning curves (magenta, right) of PV (**f**) and SST interneurons (**g**) during 30-min free exploration. Each spike is color-coded for θ_allo_ (inset, North: N; East: E; South: S; and West: W). **h**, (Left) Border score plotted as a function of egocentric vector length (VL) for all recorded PV (purple dots) and SST interneurons (green dots), with corresponding probability distributions. (Right) Box plots of VL and border scores for PV (purple) and SST interneurons (green). Vertical dotted line: 99^th^ percentile of randomly shuffled VL and border score respectively. **i**, Venn diagrams showing the proportions of egocentric boundary cells (EBCs, blue), border cells (BCs, magenta), conjunctive EBC-BC (purple), and other cell types (other) in PV (left) and SST interneurons (right). **j**, Normalized firing rate maps of EBCs aligned to preferred θ_ego_ (left) and preferred *D*_ego_ (right). **k**, Polar scatter plots of preferred θ_ego_ and preferred *D*_ego_ (left), and box plots of preferred *D*_ego_ for PV (purple) and SST interneurons (green). **l**, Normalized firing rate maps for BCs aligned to distance to the nearest environmental boundary (left), box plots of border coverage (middle), and mean firing distance to the wall (right) for PV (purple) and SST interneurons (green). Box plots (c, d, h, k, l) show median (center line), interquartile range (box), and minimum and maximum values (whiskers). Two-sided Wilcoxon rank-sum test, \*\*\**P*_Vector length_ = 1.30 × 10^-12^ (**h**), \*\*\**P*_Border score_ = 3.42 × 10^-31^ (**h**), \*\*\**P* = 8.39 × 10^-7^ (**k**), *P*_Border coverage_ = 0.463 (**l**), *P*_Mean firing distance to wall_ = 0.195 (**l**). n = 328 PV interneurons from eleven PV-Cre mice; n = 222 SST interneurons from six SST-Cre mice.

Both PV and SST interneurons exhibited spatially modulated activity (Fig. 1c,d), as analyzed by computing the correlation of firing rate maps between the first and last 15 min of free exploration. However, SST interneurons showed significantly greater map stability across time compared to PV interneurons, with a larger fraction of spatially stable cells identified in SST interneurons than PV interneurons (Fig. 1c,d). These results indicate that both populations are spatially tuned, with SST interneurons representing space more stably than PV interneurons.

Next, to investigate whether PV and SST interneurons represent environmental boundaries and their spatial reference frames, we plotted each cell’s spike-trajectory map with spikes color-coded by allocentric head direction (θ_allo_) and computed egocentric bearing (θ_ego_) and egocentric distance (*D*_ego_) to the nearest boundary (Fig. 1e). The majority of PV interneurons exhibited consistent egocentric-bearing preferences while allocentric head direction was broadly distributed (Fig. 1f and Extended Data Fig. 1a), indicating stable egocentric tuning. In contrast, SST interneurons showed less tuning for both egocentric-bearing and allocentric heading, but consistently fired near proximal boundaries (Fig. 1g and Extended Data Fig. 1b), suggesting boundary-anchored coding.

Further quantification confirmed this dissociation. We calculated egocentric vector length (VL) and border scores for each neuron (Fig. 1h). Among all recorded PV and SST interneurons, PV interneurons had significantly higher VL than SST interneurons, while SST interneurons had significantly higher border scores than PV interneurons (Fig. 1h). Neurons with VLs above the 99^th^ percentile shuffle threshold were classified as egocentric boundary cells (EBCs) and, similarly, neurons with border scores above the shuffle threshold were classified as border cells (BCs, Fig. 1i). Overall, 34 % of PV interneurons were EBCs, whereas 8 % were BCs, with only 3 % exhibiting conjunctive EBC-BC (Fig. 1i). For SST interneurons, 12 % were EBCs while 48 % were BCs, with 7 % conjunctive EBC-BC (Fig. 1i). These results reveal functional divisions between PV and SST interneurons in which PV interneurons showed a dominant egocentric bias, whereas SST interneurons favored boundary-anchored coding.

EBCs in both PV and SST interneurons covered the full egocentric bearing ranges (–180° to 180°; Fig. 1j,k). However, while PV interneurons coded egocentric boundaries at both proximal and distal distances, SST interneurons were confined to proximal ranges (Fig. 1j,k). Among neurons classified as BCs, both PV and SST interneurons exhibited large border coverage and short mean distance to the wall (Fig. 1k), indicating similar proximal boundary encoding. A minor subpopulation of PV and SST interneurons exhibited head direction tuning (Extended Data Fig. 2a-e) and conjunctive EBC-head direction tuning (Extended Data Fig. 2d).

Together, these results demonstrate that RSC PV and SST interneurons themselves encode environmental geometry using distinct spatial reference frames: PV interneurons preferentially represent egocentric coordinates, while SST interneurons encode boundary-anchored structure and stability.

### Rapid PV interneuron recruitment couples self-motion signals to egocentric coding precision

Egocentric spatial representations require continuous updating of an animal’s orientation and position during navigation. The fidelity of egocentric coding therefore depends on integrating self-motion signals such as angular head velocity (AHV) and running speed. To determine whether PV and SST interneurons differ in their coupling to these movement-related variables, we analyzed the deconvolved spikes during free exploration while tracking moment-to-moment AHV and speed.

PV interneurons exhibited strong modulation by AHV, showing sharp increases in firing rates during bidirectional, left or right head turns (Fig. 2a and Extended Data Fig. 3c). In contrast, SST interneurons were weakly modulated by AHV at the single-cell level (Fig. 2b and Extended Data Fig. 3d). AHV score (see Methods) was used in classifying neurons into positively AHV-tuned cells (Extended Data Fig. 2a), and, at the population level, a greater proportion of PV interneurons were positively tuned to AHV than SST interneurons (Fig. 2b). Overall, mean AHV tuning was significantly higher in PV than in SST interneurons (Fig. 2c, left), as quantified by the absolute slope of firing rate as a function of AHV (Fig. 2c, right). A minor subpopulation of PV and SST interneurons displayed decreased spike firing rates, but represented a small fraction of each population (Extended Data Fig. 3e,f).

**Fig. 2:**
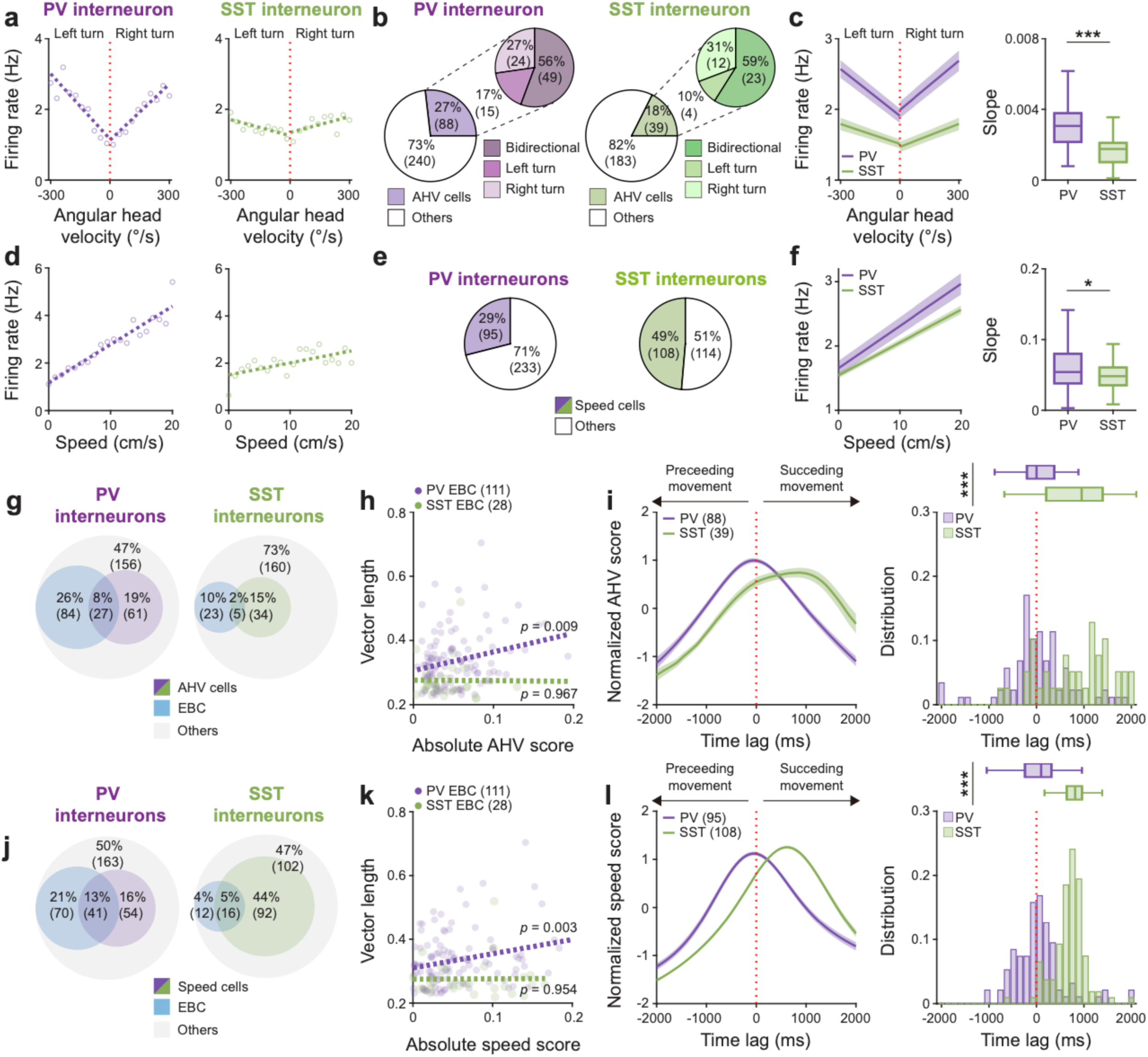
Sequential recruitment of PV and SST interneurons couples self-motion to egocentric spatial coding in the retrosplenial cortex. **a-b,** Representative angular head velocity (AHV) tuning curves of PV and SST interneurons (**a**) and pie charts showing the proportion of AHV-tuned cells in PV (purple) and SST interneurons (green) during left and right bidirectional turns (**b**). Vertical red dotted line: 0°/s. **c,** Population-averaged AHV tuning curves aligned to 0°/s (left) and box plots of AHV tuning curve slopes (right) for PV (purple) and SST interneurons (green). **d-f**, Same as **a-c,** but for locomotion speed-tuned PV and SST interneurons. **g**, Venn diagram showing the proportions of egocentric boundary cell (EBCs, blue), AHV-tuned cells (PV: purple, SST: green), and other cell types (other, gray) in PV (left) and SST interneurons (right). **h**, Egocentric boundary vector length (VL) plotted as a function of the absolute AHV score for EBC PV and SST interneurons, with linear regression fits (dotted lines) for PV (purple) and SST interneurons (green). **i,** (left) Time-shifted population mean AHV score as a function of time relative to movement onset (red dotted line: 0 ms). (right) Probability distributions of the peak time lags at which AHV coupling was maximized for PV (purple) and SST interneurons (green). **j-l,** Same as **g-i,** but for locomotion speed-tuned cells. Box plots (**c**, **f**, **i**, **l**) show median (center line), interquartile range (box), and minimum and maximum values (whiskers). Two-sided Wilcoxon rank-sum test, \*\*\**P* = 1.16 × 10^-9^ (**c**), \**P* = 0.0434 (**f**), \*\*\**P* = 6.20 × 10^-7^ (**i**), \*\*\**P* = 2.71 × 10^-20^ (**l**). n = 328 PV interneurons from eleven PV-Cre mice; n = 222 SST interneurons from six SST-Cre mice.

We next examined how neuronal activity scaled with locomotion speed. PV interneurons displayed graded, positive modulation of firing rate with increasing speed (Fig. 2d, left), while a smaller subset showed decreased responses at high speeds (Extended Data Fig. 3g,h). In contrast, SST interneurons were largely insensitive to running speed, showing weak correlations (Fig. 2d, right). At the population level, based on speed score distribution (Extended Data Fig. 3b), greater proportion of SST interneurons were tuned to speed than PV interneurons (Extended Data Fig. 2e), however, speed-tuning indices were significantly larger in PV than SST interneurons (Fig. 2f, left), as indicated by the slope of firing rates as a function of speed (Fig. 2f, right). Together, these findings suggest that PV interneurons encode self-motion signals with high gain, whereas SST interneurons contribute a distributed, lower-gain representation.

To examine the relationship between self-motion signals and egocentric tuning, we compared self-motion encoding and EBC encoding across PV and SST interneuron populations. Conjunctive AHV-EBC tuning was observed in PV interneurons, whereas AHV- and EBC-tuned SST interneurons were largely non-overlapping (Fig. 2g). Across EBC PV interneurons, AHV tuning positively correlated with egocentric VL (Fig. 2h), while no significant correlation was observed for SST interneurons (Fig. 2h). Temporal lag analysis of population activity (**see Methods**) revealed that PV interneuron activation preceded SST interneuron activation by ∼1 s during epochs of elevated AHV score (Fig. 2i). This temporal offset indicates a sequential recruitment in which self-motion signals are first engaged by PV interneurons before SST interneurons are engaged over slower timescales, suggesting a temporal division between rapid egocentric encoding and slower representational stabilization.

Similar analysis on the relationship between speed-tuned and EBC-encoding PV and SST interneuron population revealed that larger populations of PV neurons were conjunctively encoding speed and EBC, while larger populations of SST interneurons were tuned to speed alone (Fig. 2j). Consistent with this, EBC PV interneurons showed strong correlations between speed modulation and egocentric VL, while no correlation was observed in SST interneurons (Fig. 2k), but temporal sequence was preserved, with PV interneuron activation preceding SST interneuron activation (Fig. 2l). Overall, these findings indicate that PV interneurons are strongly and rapidly modulated by self-motion signals, dynamically linking movement with the precision of egocentric bearing representations, whereas SST interneurons operate on slower timescales to maintain stability.

### PV and SST interneurons exhibit distinct synchrony and population geometry for egocentric encoding

Having established that PV interneurons encode self-motion signals and are recruited earlier than SST interneurons (Fig. 2), we next examined how PV and SST interneurons are coordinated at the network level during navigation.

Spike raster visualization and pairwise synchrony analysis revealed distinct coordination motifs between the two inhibitory populations (Fig. 3a,b). PV interneurons exhibited temporally compact, clustered synchrony that was selectively enhanced among neurons sharing similar preferred θ_ego_ (Fig. 3b, top). In contrast, SST interneurons displayed broad, homogeneous synchrony across the population that was largely independent of preferred θ_ego_ and extended over longer temporal windows (Fig. 3b, bottom), showing global inhibitory coherence. Overall, spike correlations were significantly higher in SST interneurons than in PV interneurons (Fig. 3c), while spike correlations aligned by θ_ego_ were significantly higher in PV interneurons than in SST interneurons (Fig. 3d).

**Fig. 3:**
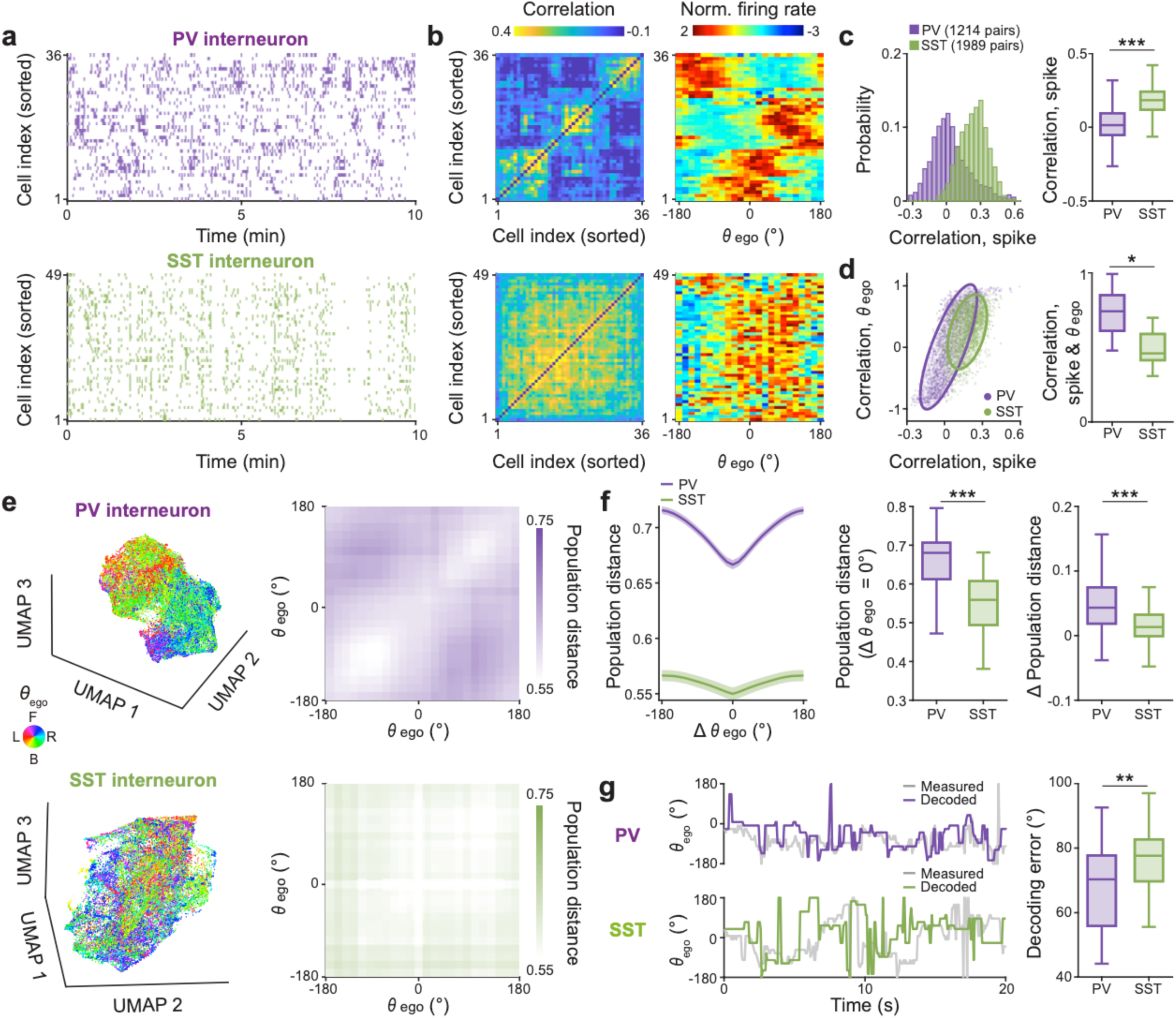
Distinct synchrony and population geometry of PV and SST interneurons in the retrosplenial cortex. **a,** Example spike raster plots of PV (purple, top) and SST interneurons (green, bottom) during 30-min free exploration in a square open chamber. Neurons are ordered based on hierarchical clustering. **b,** Population pairwise correlation similarity matrices (left) and normalized firing rate maps aligned to preferred egocentric bearing (θ_ego_) (right) for PV (top) and SST interneurons (bottom). Cells are sorted in the same order as in panel **a**. **c-d,** Probability distributions of pairwise spike correlations (**c**, left) and mean pairwise spike correlation as a function of preferred θ_ego_ (**d**, left), with corresponding box plots (right) for PV (purple) and SST interneurons (green). **e,** (left) UMAP embeddings of population activity colored by preferred egocentric bearing (θ_ego_) for PV (top) and SST interneurons (bottom). (right) Corresponding egocentric bearing-aligned population distance matrices across θ_ego_ bins. **f,** Population distance as a function of the difference in preferred egocentric bearing (Δθ_ego_, left) and box plots of population distance at Δθ_ego_ = 0° (middle) and the difference in population distance between 0° and 180° (Δ population distance; right). **g,** Bayesian decoding of θ_ego_ from PV and SST interneuron population activity. (left) Example time series of actual and decoded θ_ego_ for PV (purple) and SST interneurons (green). (right) Box plots of decoding error for PV (purple) and SST interneurons (green). Box plots (**c**, **e**, **f**) show 25th (lower box line), 50th (middle line), and 75th (upper box line) percentile values; whiskers indicate minimum and maximum values. Two-sided Wilcoxon rank-sum test, \*\*\**P* = 2.74 × 10^-274^ (**c**), \**P* = 0.0127 (**d**), \*\*\**P* _Population distance_ = 5.61 × 10^-39^ (**f**), \*\*\**P* _ΔPopulation distance_ = 2.41 × 10^-17^ (**f**), \*\**P* = 0.0093 (**g**). n = 328 PV interneurons from eleven PV-Cre mice; n = 222 SST interneurons from six SST-Cre mice.

To determine whether these distinct synchrony motifs translate into differences in population-level representation of egocentric space, we examined the structure of inhibitory population activity using nonlinear dimensionality reduction. UMAP embedding of population activity revealed that PV interneurons formed structured clusters organized by egocentric bearing, with neurons preferring similar θ_ego_ occupying nearby positions in low-dimensional space (Fig. 3e and Extended Data Fig. 4a). In contrast, SST interneuron population activity was more diffusely distributed, with no clear clustering by egocentric bearing (Fig. 3e and Extended Data Fig. 4b). We quantified this organization by computing pairwise population distances as a function of θ_ego_ difference. In PV interneurons, population distance increased systematically with increasing θ_ego_ separation, indicating that egocentric bearing similarity is reflected in population geometry (Fig. 3f). This relationship was absent in SST interneurons, where population distances were largely independent of θ_ego_ difference (Fig. 3f). Across sessions, the difference in population distance between similar and dissimilar bearings was significantly larger for PV than for SST interneurons (Fig. 3f). We then asked whether these population-level differences affect the information content of inhibitory activity. Using a Bayesian decoder (**see Methods**), we decoded θ_ego_ from PV and SST population activity. Decoding accuracy was significantly higher—and decoding error significantly lower—for PV interneurons than for SST interneurons (Fig. 3g), indicating that PV population activity carries more precise information about egocentric orientation.

Together, these results show that PV and SST interneurons differ not only in their synchrony structure, but also in how egocentric space is embedded in population activity. PV interneurons form bearing-aligned, information-rich population states with structured geometry, whereas SST interneurons exhibit globally coherent but egocentrically unstructured population dynamics.

### PV and SST inhibition differentially disrupt egocentric coding of space by retrosplenial excitatory neurons

The distinct spatial reference frames (Fig. 1), self-motion coupling (Fig. 2), and inhibitory synchrony motifs (Fig. 3) observed in PV and SST interneurons predict complementary causal influences on egocentric representations in RSC excitatory (EX) neurons. To directly test how these distinct inhibitory dynamics shape excitatory neuronal representations of egocentric space, we performed Ca^2+^ imaging on GCaMP6s-expressing RSC EX neurons while optogenetically silencing either halorhodopsin-expressing PV or SST interneurons (Fig. 4a,b). Optogenetic silencing of PV or SST interneurons slightly reduced the overall firing rate of RSC excitatory neurons (Fig. 4c), indicating modest changes in baseline activity without gross disruption of network dynamics.

**Fig. 4:**
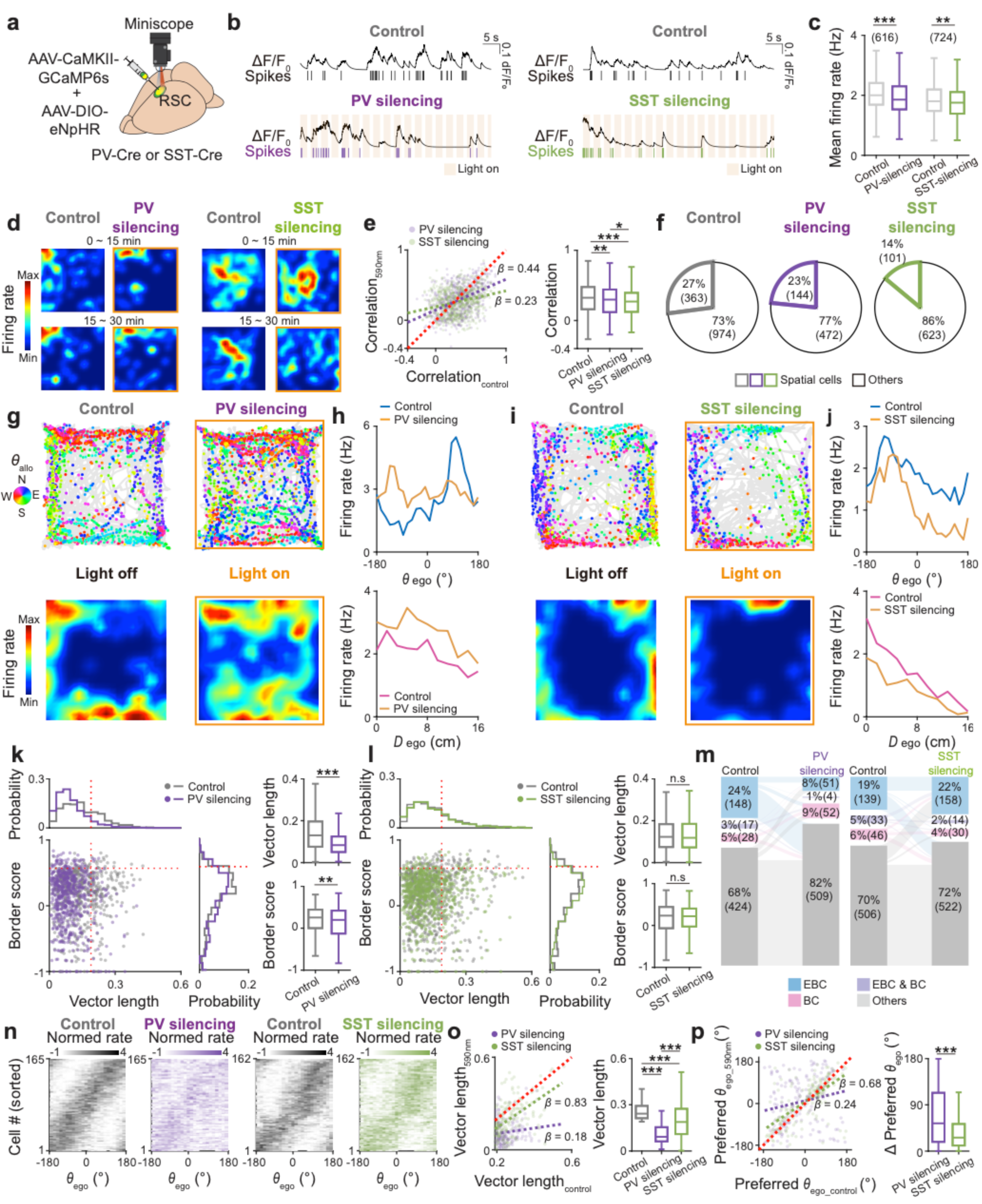
Distinct effects of PV and SST interneuron silencing on egocentric spatial coding of RSC excitatory neurons in the retrosplenial cortex. **a,** Schematic of experimental set up for AAV-CaMKII-GCaMP6s and AAV-DIO-eNpHR injections, GRIN lens implantation, and miniaturized microscope-based Ca² imaging of retrosplenial cortex (RSC) excitatory neurons during optogenetic silencing of NpHR-expressing PV or SST interneurons using orange light (590-650 nm) in PV-Cre and SST-Cre mice. **b,** Representative raw Ca² signals with corresponding deconvolved spikes of RSC excitatory neurons under the control condition (black, top) and during optogenetic silencing of NpHR-expressing PV (purple, bottom left) and SST interneurons (green, bottom right). **c,** Box plots of mean firing rates of RSC excitatory neurons under the control conditions and during optogenetic silencing of NpHR-expressing PV (purple) or SST interneurons (green). **d,** Representative spatial firing rate maps of RSC excitatory neurons during the first (0 – 15 min) and second halves (15 – 30 min) of 30-min free exploration in a square open chamber under control conditions (left) and during optogenetic silencing (right, orange square border) of NpHR-expressing PV (left) and SST interneurons (right). **e-f,** Scatter plots of spatial stability correlations between spatial firing rate maps from the first and last 15-min epochs (**e**) under the control conditions and during optogenetic silencing of NpHR-expressing PV (purple) and SST interneurons (green) (left), with corresponding box plots (middle). Pie charts (**f**) showing the proportions of spatially selective and non-spatially selective cells under the control conditions (gray) and during optogenetic silencing of NpHR-expressing PV (purple) and SST interneurons (green). **g,** Representative spike-trajectory plots (mouse trajectory: gray line, spike locations: colored dots) under control (top, left) and during optogenetic silencing of NpHR-expressing PV interneurons (top, right), with corresponding firing rate maps (bottom).**h,** (Top) Egocentric bearing (θ_ego_) tuning curves of RSC excitatory neurons under control conditions (dark blue) and during optogenetic silencing of NpHR-expressing PV interneurons (orange). Each spike is color-coded by θ_allo_ (inset, North: N; East: E; South: S; and West: W). (bottom) Egocentric distance (*D*_ego_) tuning curves under control conditions (magenta) and during optogenetic silencing of NpHR-expressing PV interneurons (orange). **i-j,** Same as **g-h,** but during optogenetic silencing of NpHR-expressing SST interneurons. **k,** (Left) Border score plotted as a function of egocentric vector length (VL) for all recorded excitatory neurons under control conditions (gray dots) and during optogenetic silencing of PV interneurons (purple dots), with corresponding probability distributions (top, right). (right) Box plots of VL (top) and border score (bottom). Vertical red dotted line: 99^th^ percentile of randomly shuffled distribution of VL and border score respectively. **l,** Same as in **k,** but during optogenetic silencing of SST interneurons. **m,** Sankey diagrams showing cell classification transitions of egocentric boundary cells (EBCs, blue), border cells (BCs, magenta), conjunctive EBC-BC (purple), and other cell types (white) between the control conditions and optogenetic silencing of NpHR-expressing PV (purple) and SST interneurons (green). **n,** Normalized egocentric bearing (θ_ego_) tuning maps of EBCs under control conditions and during optogenetic silencing of NpHR-expressing PV and SST interneurons. Cells are ordered by preferred θ_ego_ in the control condition. **o,** (Left) VL during optogenetic silencing of PV (purple) and SST interneurons (green) plotted as a function of VL under the control conditions, with linear regression fits (dotted line, inset: slope (β coefficient)). (Right) Corresponding box plots of VL. **p,** (Left) θ_ego_ under optogenetic silencing of NpHR-expressing PV (purple) or SST interneurons (green) plotted as a function of that under the control condition with linear regression fit (dotted line, inset: slope (β coefficient)). (right) Corresponding changes in θ_ego_ (right). Box plots (**c**, **e**, **k**, **l**, **o**, **p**) show median (center line), interquartile range (box), and minimum and maximum values (whiskers).. Paired t test, \*\*\**P*_Control-PV silencing_ = 1.57 × 10^-6^ (**c**), \*\**P*_Control-SST silencing_ = 0.0030 (**c**), \*\*\**P*_Vector length_ = 1.99 × 10^-30^ (**k**), \*\*\**P*_Border score_ = 7.42 × 10^-4^ (**k**), *P*_Vector length_ = 0.5970 (**l**), *P*_Border score_ = 0.7642 (**l**). Two-sided Wilcoxon rank-sum test, \*\**P*_Control-PV silencing_ = (**e**), \*\*\**P*_Control-SST silencing_ = 4.16 × 10^-9^ (**e**), \*\*\**P*_PV silencing-SST silencing_ = 0.0212 (**e**), \*\*\**P*_Control-PV silencing_ = 9.00 × 10^-54^ (**o**), \*\*\**P*_Control-SST silencing_ = 1.35 × 10^-12^ (**o**), \*\*\**P*_PV silencing-SST silencing_ = 3.64 × 10^-34^ (**o**), \*\*\**P* = 9.73 × 10^-6^ (**p**). n = 616 excitatory neurons from seven PV-Cre mice; n = 724 excitatory neurons from eight SST-Cre mice.

We first examined the effect of inhibitory perturbation on spatial stability of EX neurons (Fig. 4d). Under control conditions, EX neurons exhibited stable spatial firing, as quantified by correlations between firing rate maps from early (0-15 min) to late exploration (15-30 min) epochs (Fig. 4d,e). PV interneuron silencing produced a modest reduction in spatial stability (Fig. 4d,e) and spatially selective neural population (Fig. 4f). In contrast, SST interneuron silencing significantly reduced spatial correlations (Fig. 4d,e) and reduced the fraction of spatially selective neurons (Fig. 4f), indicating that long-timescale stability of excitatory spatial representations depends preferentially on SST inhibition.

We next asked how PV and SST interneurons modulate egocentric spatial tuning of EX neurons. Under control conditions, EX neurons displayed robust egocentric boundary tuning, characterized by preferred θ_ego_ and *D*_ego_ relative to the nearest wall (Fig. 4g). However, silencing PV interneurons led to a marked disruption of egocentric bearing selectivity and egocentric distance coding; spike-trajectory and firing rate maps became disorganized (Fig. 4g,h, and Extended Data Fig. 5a). In contrast, silencing SST interneurons largely preserved θ_ego_ selectivity in EX neurons while boundary-anchored firing was slightly weakened (Fig. 4i,j and Extended Data Fig. 5b).

Quantification of egocentric VL and border score of EX confirmed these dissociable effects: PV interneuron silencing significantly reduced both VL and border score (Fig. 4k), whereas SST silencing had no effect on either (Fig. 4l). Accordingly, the fraction of EBC EX neurons decreased following PV interneuron silencing, but was unchanged by SST interneuron silencing, while the fraction of BCs remained similar across conditions (Fig. 4m). To further characterize how PV and SST interneurons modulate EX EBCs, we analyzed the VL and preferred θ_ego_ before and after PV or SST interneuron silencing (Fig. 4n-p). Significant reductions in VL and shifts in preferred θ_ego_were induced by PV interneuron silencing but not by SST interneuron silencing (Fig. 4n-p). A subpopulation of RSC EX neurons exhibited head direction tuning, but was minimally affected by either PV or SST interneuron silencing (Extended Data Fig. 6).

Together, these results demonstrate dissociable roles of PV and SST interneurons in egocentric coding in RSC EX neurons: PV inhibition sharpens egocentric orientation precision, whereas SST inhibition supports stabilization of environmental geometry.

### PV interneurons couple self-motion integration signals to egocentric coding precision in RSC excitatory neurons

Distinct inhibitory dynamics in PV and SST interneurons suggest that they may differentially control how self-motion signals are integrated into excitatory spatial representations. To test this directly, we examined how optogenetic silencing of PV or SST interneurons alters self-motion modulation and egocentric tuning in RSC EX neurons. Under control conditions, RSC EX neurons displayed robust modulation by AHV, with activity increasing during head turns (Fig. 5a). Optogenetic silencing of PV interneurons markedly reduced AHV modulation across the EX population, indicating a loss of coupling between rotational self-motion and excitatory activity (Fig. 5a). In contrast, silencing SST interneurons produced minimal changes in AHV modulation, despite strongly disrupting spatial stability and boundary anchoring (Fig. 5a). Using AHV score to classify positively AHV-tuned cells, both PV and SST interneuron silencing reduced the proportion of AHV-tuned EX neurons (Fig. 5b). However, AHV tuning strength was significantly decreased by PV interneuron silencing than by SST interneuron silencing (Fig. 5c), as quantified by the slope of firing rate as a function of AHV (Fig. 5d).

**Fig. 5:**
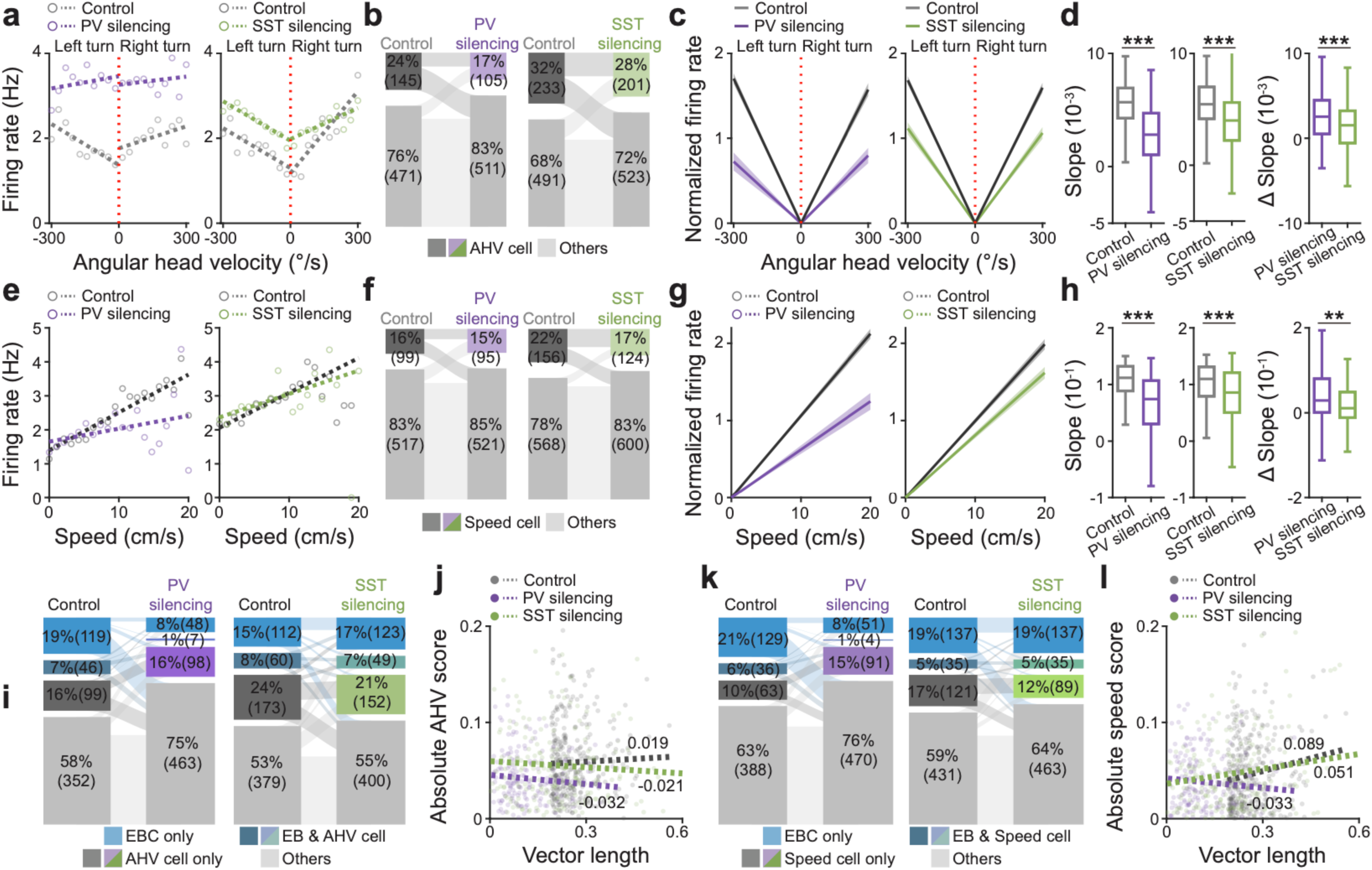
Dissociable roles of PV and SST interneurons in self-motion tuning and egocentric coding precision in the retrosplenial cortex. **a,** Representative angular head velocity (AHV) tuning curves of retrosplenial cortex (RSC) excitatory neurons under control conditions (gray) and during optogenetic silencing of NpHR-expressing PV (purple) and SST interneurons (green). Vertical red dotted line: 0°/s. **b-d,** Sankey diagrams of AHV cell classification transitions (**b**), population-averaged AHV tuning curves aligned to 0°/s (**c**), box plots of AHV tuning curve slopes (**d**, left) and the mean changes in slope (**d**, right) under the control conditions (gray) and during optogenetic silencing of NpHR-expressing PV (purple) and SST interneurons (green). **e-h,** Same as **a-d,** but for locomotion speed-tuned cells. **i,** Sankey diagrams showing cell classification transitions between egocentric boundary cells (EBCs, blue) AHV-tuned cells (control: cyan; PV silencing: purple; SST silencing: green), and conjunctive EBC–AHV cells (dark blue) under the control conditions and during optogenetic silencing of PV or SST interneurons. **j,** Absolute AHV score plotted as a function of egocentric boundary vector length (VL) for EBC excitatory neurons under the control conditions (gray dots) and during optogenetic silencing of PV (purple) and SST interneurons (green) with linear regression fits (dotted lines, inset: slope (β coefficient)). **k-l,** Same as **i-j,** but for locomotion speed-tuned cells. Box plots (**d**, **h**) show median (center line), interquartile range (box), and minimum and maximum values (whiskers). Paired t test, \*\*\**P*_Control-PV silencing_ = 1.11 × 10^-18^ (**d**), \*\*\**P*_Control-SST silencing_ = 1.69 × 10^-12^ (**d**), \*\*\**P*_Control-PV silencing_ = 9.33 × 10^-10^ (**h**), \*\**P*_Control-SST silencing_ = 4.27 × 10^-6^ (**h**). Two-sided Wilcoxon rank-sum test, \*\*\**P*_PV silencing-SST silencing_ = 7.39 × 10^-4^ (**d**), \*\*\**P*_PV silencing-SST silencing_ = 0.0031 (**h**). n = 616 neurons from 7 PV-Cre mice; n = 724 neurons from 8 SST-Cre mice.

We next examined modulation of EX neurons by locomotion speed. Under control conditions, EX activity scaled with running speed, with higher firing rates during fast locomotion epochs (Fig. 5e). PV interneuron silencing significantly reduced speed modulation indices across the EX population, whereas SST interneuron silencing had little effect on speed tuning (Fig. 5e). At the population level, SST interneuron silencing decreased the proportion of EX neurons tuned to speed, while PV interneuron silencing had minimal effect (Fig. 5f). Nonetheless, speed-tuning indices were significantly decreased by PV than SST interneuron silencing (Fig. 5g), as indicated by the slope of firing rate as a function of speed (Fig. 5h).

To examine how inhibitory perturbations alter the relationship between self-motion signals and egocentric tuning in RSC EX neurons, we compared changes in self-motion encoding and EBC encoding across perturbation conditions (Fig. 5i-l). Both EBC and conjunctive AHV-EBC tuning in EX neurons were significantly reduced by PV interneuron silencing, while SST interneuron silencing increased both EBC and conjunctive AHV-EBC populations (Fig. 5i). Directly comparing the effect on VL, the positive correlation between AHV tuning and egocentric VL in EX neurons was markedly reduced by silencing PV interneurons, abolishing the AHV-VL relationship (Fig. 5j). Silencing SST interneurons produced minimal changes in AHV modulation and preserved the AHV–VL relationship (Fig. 5j). Similar results were observed for speed-EBC relationships (Fig. 5k-l). Together, these results indicate that PV-mediated inhibition is required to maintain the coupling between self-motion signals and the precision of egocentric tuning in excitatory neurons, whereas SST-mediated inhibition has minimal effect on self-motion coupling, consistent with a role in integrating the output of PV encoding rather than gating self-motion signals directly.

### PV and SST interneurons differentially regulate the population geometry of egocentric boundary cells

To determine how PV and SST interneurons shape the geometry of egocentric representations in RSC excitatory neurons, we quantified population structure and decoding performance during optogenetic silencing of each subtype.

We first examined the organization of excitatory population states using principal component analysis (PCA). Under control conditions, EX EBCs formed discrete clusters in low-dimensional space that were organized according to preferred θ_ego_ (Fig. 6a), indicating structured population states encoding egocentric orientation. During PV interneuron silencing, the overall cluster topology and bearing-dependent grouping were preserved (Fig. 6b). However, correlations among neuron pairs with similar egocentric bearings were significantly reduced (Fig. 6b,c), particularly between neuron pairs with aligned (0°) and opposing (180°) egocentric bearings, indicating degradation of egocentric state precision despite preservation of the global cluster structure. In contrast, SST interneuron silencing disrupted the overall organization in low-dimensional space (Fig. 6b). Cluster separability was markedly reduced and correlations across θ_ego_ were broadly disrupted (Fig. 6b,d), indicating a collapse of structured egocentric population organization.

**Fig. 6:**
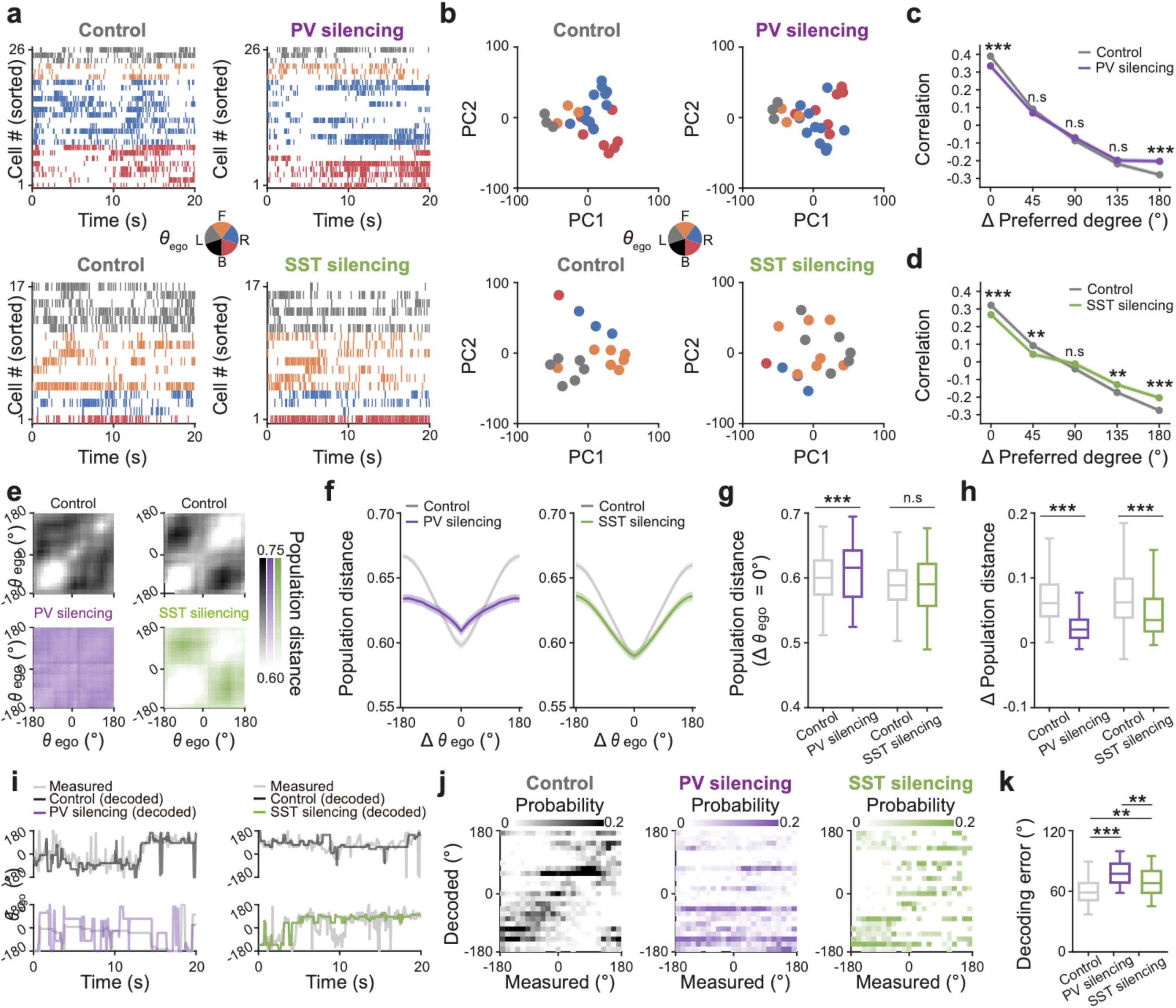
Distinct roles of PV and SST interneurons in shaping excitatory population geometry. **a,** Representative spike raster plots of RSC excitatory egocentric boundary cells (EBCs) recorded from PV-Cre mice (top) and SST-Cre mice (bottom) under the control conditions (control, left) and during optogenetic silencing of NpHR-expressing PV or SST interneurons using 590 nm light (right). Each spike is color-coded by preferred egocentric bearing (θ_ego_) (inset, Front: F; Left: L; Back: B; and Right: R). **b,** Corresponding population activity EBC excitatory neurons projected onto the first two principal components (PC1 vs PC2), with each point representing a single time bin, colored by preferred θ_ego_. (inset). **c-d,** Mean pairwise correlations between EBCs plotted as a function of the difference in preferred θ_ego_ (Δ preferred θ_ego_) under the condition and during optogenetic silencing of PV (**c**, purple) and SST interneurons (**d**, green). **e-h,** Population distance matrices across θ_ego_ of RSC excitatory EBCs (**e**), population distance as a function of the difference in θ_ego_ (Δθ_ego_) (**f**), box plots of population distance aligned to 0° (**g**) and the difference in population distance at 0° and 180° (Δ population distance; **h**) under the control (gray) and during optogenetic silencing of NpHR-expressing PV interneurons (purple) and SST interneurons (green). **i-k,** Representative time series of actual and decoded θ_ego_ (**i**) from RSC excitatory EBCs, confusion matrices of decoding performance (**j**) and box plots of mean decoding error (**k**) under control (gray) and during optogenetic silencing of NpHR-expressing PV interneurons (purple) and SST interneurons (green). Box plots (**g**, **h**, **k**) show 25th (lower box line), 50th (middle line), and 75th (upper box line) percentile values; whiskers indicate minimum and maximum values. Paired t test, \*\*\**P*_0_ = 6.09 × 10^-6^ (**c**), *P*_45_ = 0.1686 (**c**), *P*_90_ = 0.2476 (**c**), *P*_135_ = 0.1869 (**c**), \*\*\**P*_180_ = 2.13 × 10^-6^ (**c**), \*\*\**P*_0_ = 2.40 × 10^-7^ (**d**), \*\**P*_45_ = 0.0016 (**d**), *P*_90_ = 0.0914 (**d**), \*\**P*_135_ = 0.0024 (**d**), \*\*\**P*_180_ = 1.59 × 10^-7^ (**d**), \*\*\**P*_Control-PV silencing_ = 5.90 × 10^-4^ (**g**), *P*_Control-SST silencing_ = 0.8035 (**g**), \*\*\**P*_Control-PV silencing_ = 1.60 × 10^-31^ (**h**), \*\*\**P*_Control-SST silencing_ = 1.08 × 10^-18^ (**h**). Two-sided Wilcoxon rank-sum test, \*\*\**P*_Control-PV silencing_ = 9.62 × 10^-19^ (**k**), \*\**P*_Control-SST silencing_ = 0.0021 (**k**), \*\**P*_PV silencing-SST silencing_ = 0.0047 (**k**). n = 616 neurons from seven PV-Cre mice; n = 724 neurons from eight SST-Cre mice.

We next quantified population geometry by computing pairwise population distances across egocentric bearing differences. Under control conditions, population distance increased systematically with increasing differences in preferred θ_ego_ (Fig. 6d,e), reflecting structured representation of egocentric space. PV interneuron silencing selectively reduced this structured distance relationship (Fig. 6f,g), consistent with reduced precision of egocentric population states. In contrast, SST interneuron silencing broadly flattened the population distance structure across all θ_ego_ differences (Fig. 6d-g), indicating disruption of the global geometry of egocentric representations.

Finally, we asked whether these geometry changes altered the information content of excitatory population activity. Under control conditions, Bayesian decoding accurately reconstructed θ_ego_ from EX EBC population activity (Fig. 6h,i). PV interneuron silencing produced a larger increase in decoding error in θ_ego_ (Fig. 6i,j). PV interneuron silencing produced a larger increase in decoding error while preserving coarse bearing discrimination, consistent with reduced precision of egocentric coding. In contrast, SST interneuron silencing resulted in more diffuse and blurred confusion matrices, indicating disruption of structured egocentric representations despite a smaller increase in decoding error.

Together, these results demonstrate that PV and SST interneurons differentially regulate the geometry of excitatory population activity in RSC: PV inhibition sharpens the precision of egocentric population states, whereas SST inhibition preserves the global organization and stability of these representations across egocentric bearings.

### PV and SST interneurons dissociate egocentric orientation precision and stability during navigation

We next asked whether the dissociable effects of PV and SST interneuron silencing on EX population organization translated into behavioral deficits during navigation. For this, we bilaterally injected AAV-Ef1a-DIO-NpHR-eYFP into RSC and implanted optic fibers (Fig. 7a) to optogenetically silence NpHR-expressing PV or SST interneurons (Fig. 7b) during a barrier-detour, goal-directed navigation task in which mice learned to navigate toward a goal through a narrow opening (Fig. 7c). In this task, θ_ego_ was defined as the egocentric bearing from the animal’s current position toward the barrier gap end, computed relative to the animal’s instantaneous heading direction (Fig. 7c).

**Fig. 7:**
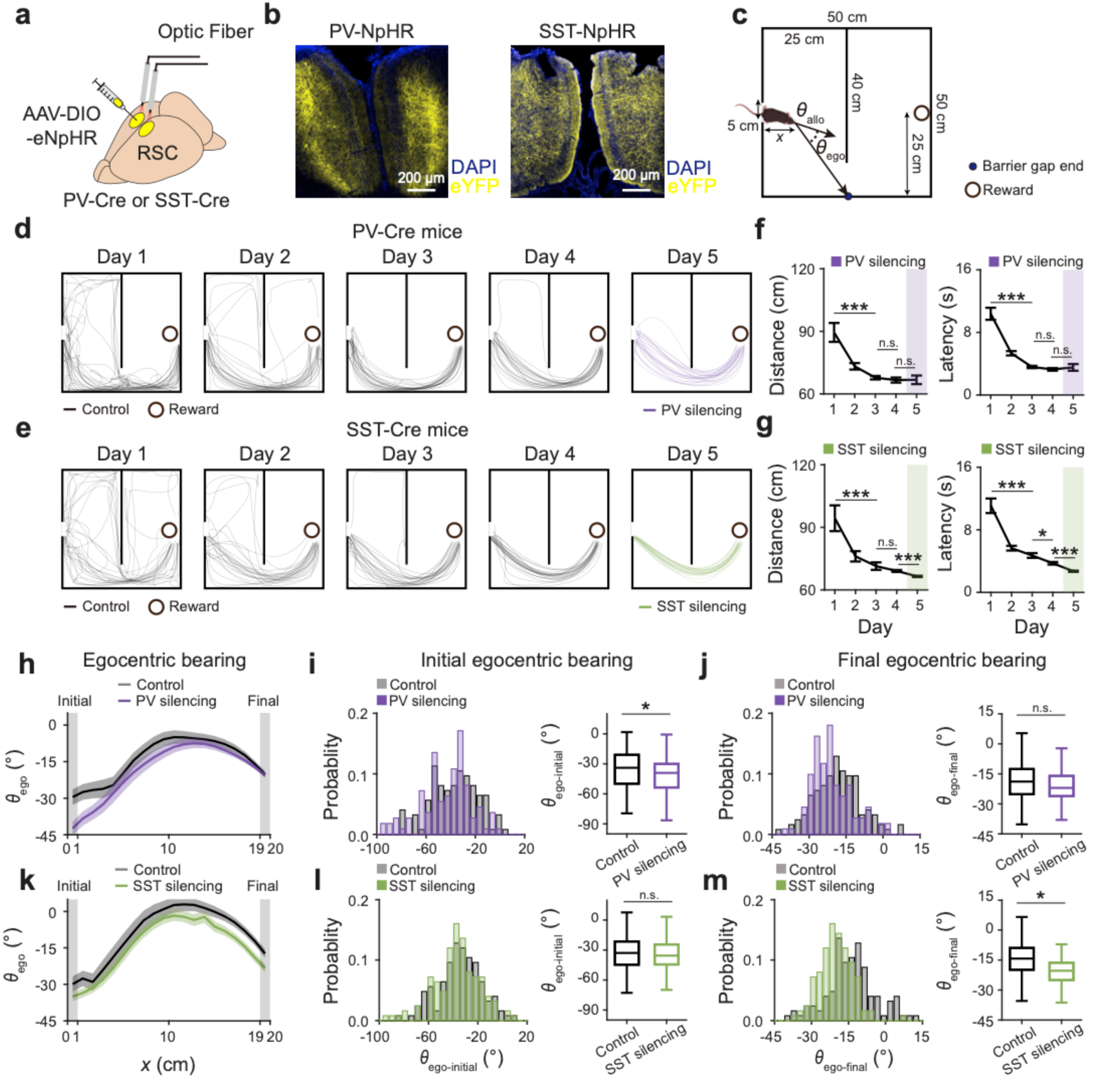
Double dissociation of PV and SST interneurons in guiding goal-directed spatial navigation. **a,** Schematic of the experimental setup for AAV-DIO-eNpHR injection and bilateral optic fiber implantation for optogenetic silencing of NpHR-expressing PV or SST interneurons in the retrosplenial cortex (RSC) during a barrier-detour goal-directed navigation task using 595 nm light in PV-Cre and SST-Cre mice. **b,** Representative fluorescence images showing optic fiber track and NpHR-expressing PV interneurons (left) and SST interneurons (right). **c**, Illustration of the barrier-detour goal-directed navigation task chamber showing distance from the start wall (*x*), the barrier gap end (blue dot), the reward location (black open circle), allocentric head direction (θ_allo_) and egocentric bearing to the barrier gap end (θ_ego_) relative to the animal’s instantaneous heading direction. **d-e,** Representative mouse trajectories from the start to reward (black circle) across Days 1-4 (gray lines) and during optogenetic silencing of NpHR-expressing PV interneurons (purple lines) on Day 5 (**d**), and the corresponding mean trajectory distance (left) and latency (right) from the start box to the reward location (**e**). **f-g,** Same as **d**,**e** but for optogenetic silencing of NpHR-expressing SST interneurons (green lines) on Day 5. **h**, θ_ego_ toward the barrier gap end as a function of *x* during the control condition (Day 4, black) and during PV interneuron silencing (purple) on Day 5. Solid lines indicate the mean and shaded areas represent mean ± SEM. Gray shaded regions indicate the first and the last spatial bin. **i,** (left) Probability distribution of θ_ego-initial_, defined as the mean θ_ego_ within the first spatial bin (*x* = 0–1 cm), comparing control and PV interneuron silencing conditions. (right) Box plots of θ_ego-initial_ comparing control and PV interneuron silencing conditions. **j**. Same as **i**, but for θ_ego-final_, defined as the mean θ_ego_ within the last spatial bin (*x* = 19–20 cm). **k-m,** Same as **h-j,** but during optogenetic silencing of SST interneurons. Box plots (**g**, **i**) show 25th (lower box line), 50th (middle line), and 75th (upper box line) percentile values; whiskers indicate minimum and maximum values. Student’s t test, \*\*\**P*_Day 1–Day 3_ = 6.82 × 10^-6^ (**f**, left), *P*_Day 3–Day 4_ = 0.3310 (**f**, left), *P*_Day 4–Day 5_ = 0.5923 (**f**, left), \*\*\**P*_Day 1–Day 3_ = 7.77 × 10^-16^ (**f**, right), *P*_Day 3–Day 4_ = 0.1765 (**f**, right), *P*_Day 4–Day 5_ = 0.6615 (**f**, right), \*\*\**P*_Day 1–Day 3_ = 3.42 × 10^-12^ (**g**, left), *P*_Day 3–Day 4_ = 0.6497 (**g**, left), \*\*\**P*_Day 4–Day 5_ = 1.37 × 10^-5^ (**g,** left), \*\*\**P*_Day 1–Day 3_ = 1.42 × 10^-14^ (**g**, right), \**P*_Day 3–Day 4_ = 0.018 (**g**, right), \*\*\**P*_Day 4–Day 5_ = 1.08 × 10^-7^ (**g**, right). Watson-Wiliams test, \**P*_Control-PV silencing_ = 0.0377 (**i**), *P*_Control-PV silencing_ = 0.5296 (**j**), *P*_Control-SST silencing_ = 0.6095 (**l**), \**P*_Control-SST silencing_ = 0.0165 (**m**). n = 113 trajectories from five PV-Cre mice; n = 125 trajectories from five SST-Cre mice.

Up to twenty-five trials of the barrier-detour task were performed each day over four consecutive days (Days 1 – 4), with PV or SST interneurons optogenetically silenced on Day 5 (Fig. 7d,e). Under control conditions, trajectory distance and latency decreased significantly from Day 1 to Day 4, indicating successful task learning (Fig. 7f,g). Optogenetic silencing of PV interneurons had no significant effect on trajectory distance or latency (Fig. 7f), indicating that optogenetic silencing did not affect overall task performance *per se*. However, optogenetic silencing of SST interneurons significantly reduced trajectory distance and latency compared to those on Day 4 (Fig. 7g).

Analysis of θ_ego_ toward the barrier gap end revealed a double dissociation between PV and SST interneuron silencing (Fig. 7h-m). During PV interneuron silencing, θ_ego_ at navigation onset was significantly reduced relative to control, indicating impaired initial egocentric orientation toward the barrier gap end (Fig. 7h,i). However, θ_ego_ progressively recovered over the course of navigation, converging toward control levels by trial end (Fig. 7h,j). In contrast, during SST interneuron silencing, initial θ_ego_ was intact and comparable to control, but θ_ego_ failed to reach control levels by the end of navigation, indicating a selective disruption of egocentric orientation maintenance over the course of navigation (Fig. 7k-m). Analysis of θ_ego_ toward the reward location during the post-barrier approach revealed no significant effect of either PV or SST interneuron silencing (Extended Data Fig. 7), consistent with the post-barrier trajectory being primarily visually guided once the reward location becomes directly visible following barrier passage.

Together, these results demonstrate that PV interneurons are required for the precise initial determination of egocentric orientation at navigation onset, while SST interneurons are required for the sustained update and maintenance of egocentric bearing during ongoing navigation. These behavioral dissociations directly mirror the sequential encoding-integration architecture identified at the circuit level, linking PV and SST inhibitory mechanisms to distinct aspects of goal-directed spatial navigation.

## Discussion

Our study identifies a division-of-labor inhibitory architecture in the RSC that links self-motion integration to the precision and stability of egocentric spatial representations. Using cell-type–specific imaging, optogenetic perturbations, and population-level analyses during navigation, we demonstrate that PV and SST interneurons make dissociable contributions to egocentric coding (Fig. 1–6). PV interneurons provide fast, self-motion–coupled inhibition that sharpens egocentric orientation and population-state precision, whereas SST interneurons stabilize boundary-anchored representations and preserve global population organization over time (Fig. 1–6). Consistent with these circuit and population-level mechanisms, perturbing these inhibitory populations produced distinct behavioral deficits during navigation (Fig. 7), with PV interneuron silencing impairing initial egocentric orientation determination, and SST interneuron silencing impairing its sustained update during ongoing navigation. Together, these findings identify inhibitory microcircuits as a key mechanism by which RSC continuously updates egocentric spatial representations while maintaining stable navigation.

A central finding of this study is that inhibitory interneurons in RSC are themselves segregated by spatial reference frame (Fig. 1). PV interneurons preferentially encoded egocentric boundary orientation and exhibited broad coverage across egocentric bearing and distance, whereas SST interneurons preferentially encoded boundary proximity and displayed greater spatial stability across time. Rather than simply modulating excitatory spatial codes, these observations indicate that reference-frame specialization is embedded within inhibitory microcircuits. Previous work has shown that excitatory neurons in RSC encode egocentric boundaries, environmental vertices, and heading-relative spatial geometry ^13–17, 34, 35^, supporting the view that RSC contributes to transformations between egocentric and allocentric spatial representations ^5, 6, 11^. Our findings extend this framework by demonstrating that inhibitory interneurons themselves participate directly in these spatial representations (Fig. 1). This organization contrasts with hippocampal and entorhinal circuits, where inhibitory interneurons primarily regulate spatial representations generated by excitatory neurons ^25–27^.

Egocentric navigation critically depends on accurate integration of self-motion signals conveying moment-to-moment changes in orientation ^36–38^. PV interneurons were strongly modulated by AHV and locomotion speed and frequently exhibited conjunctive tuning with EBC (Fig. 2). Moreover, PV interneuron activity preceded SST interneuron activation during periods of elevated self-motion, revealing a temporal sequence in inhibitory engagement (Fig. 2). At the network level, PV interneurons exhibited spatially structured synchrony that was strongest among neurons sharing similar egocentric bearing preferences (Fig. 3). In contrast, SST interneurons displayed slower and more globally coherent synchrony that was largely independent of egocentric bearing (Fig. 3). These distinct synchrony motifs suggest that PV interneurons coordinate spatially specific inhibitory ensembles that sharpen egocentric population states, whereas SST interneurons provide distributed inhibitory coherence across the network ^39^. Such functional specialization parallels the broader division of labor between fast perisomatic PV inhibition and dendritic SST inhibition observed across cortical circuits ^21, 22, 28, 29^. The sequential recruitment of PV before SST interneurons is consistent with short-term synaptic plasticity profiles, whereby PV interneurons are rapidly engaged via depressing synapses while SST interneurons require facilitated inputs for activation ^40^. This temporal sequence may reflect a functional division between encoding and integration of egocentric spatial information: PV interneurons rapidly encode moment-to-moment changes in egocentric orientation with high temporal precision, while SST interneurons — activated subsequently over slower timescales—integrate the output of this encoding to stabilize the resulting spatial representation. Importantly, SST interneurons appear to integrate not the self-motion signals themselves, but their encoded consequences—consistent with the finding that SST silencing does not impair self-motion modulation of excitatory neurons (Fig. 5), while strongly disrupting long-timescale spatial stability (Fig. 4). Notably, PV interneurons exhibit stronger phase-locking and fire at earlier phases than SST interneurons during cortical network oscillation ^39^, consistent with their preferential contribution to feedforward over feedback inhibition in cortical circuits ^41^, similar to the bearing-aligned synchrony and earlier recruitment of PV interneurons we observed in RSC during navigation (Fig. 3).

The functional consequences of these inhibitory dynamics were revealed by optogenetic perturbation experiments. Silencing PV interneurons degraded egocentric tuning in EX neurons, reduced decoding accuracy of egocentric bearing, and disrupted the coupling between self-motion signals and egocentric coding precision (Fig. 4–6). In contrast, perturbing SST interneurons destabilized spatial representations over longer timescales, reducing spatial stability of EX neurons, weakening boundary anchoring, and disrupting the global organization of egocentric population states (Fig. 4–6). At the population level, PV inhibition preserved the overall structure of egocentric population states but increased dispersion within these states, indicating a selective loss of representational precision rather than a collapse of spatial organization (Fig. 6). In contrast, SST interneuron inhibition disrupted the global organization and stability of these states (Fig. 6). Similar stabilization roles have been described for dendrite-targeting SST interneurons in hippocampal circuits, where they regulate long-timescale stability of place cell representations and persistent spatial coding ^26, 28, 42^, though their synaptic mechanism in RSC remains to be established. These results provide experimental evidence that inhibitory circuits regulate the geometry of neural population activity rather than simply modulating firing rate or tuning ^43, 44^. Theoretical work has suggested that spatial representations emerge from continuous network dynamics that combine velocity-driven updating with stabilizing constraints ^20, 44–46^. Our results suggest that separable inhibitory circuits may implement these dissociable mechanisms.

Finally, the circuit mechanisms identified here translated directly into behavioral consequences during navigation (Fig. 7). PV interneuron silencing selectively impaired initial egocentric orientation at navigation onset—consistent with a role in rapidly encoding self-referenced bearing—but θ_ego_ progressively recovered toward control levels over the course of navigation, suggesting that alternative mechanisms can compensate over time in the absence of PV-mediated encoding. In contrast, SST interneuron silencing preserved initial θ_ego_ but produced a selective failure to sustain egocentric orientation maintenance toward the goal, with θ_ego_ diverging from control levels as navigation progressed—consistent with a role in the sustained integration and maintenance of egocentric bearing rather than its initial encoding. These behavioral dissociations directly mirror the sequential encoding-integration architecture identified at the circuit level. Whether these dissociable inhibitory mechanisms generalize to other navigational contexts beyond the barrier-detour task remains an important direction for future work.

Several limitations of the present study merit consideration. First, the temporal resolution of calcium imaging constrains interpretation of the ∼1 s PV-SST lag, which reflects population-level dynamics rather than direct synaptic connectivity. Second, halorhodopsin-mediated silencing may produce rebound excitation following light offset, which could contribute to some of the observed effects. Third, the circuit mechanisms by which PV interneurons receive self-motion inputs to encode egocentric orientation—and by which SST interneurons generate globally coherent synchrony to stabilize spatial representations—remain unknown. Whether PV interneurons receive direct thalamic or subcortical self-motion signals, and what drives the globally coherent dynamics of SST interneurons in RSC, are important questions for future investigation. Additionally, cortical SST interneurons are known to provide inhibitory input onto PV interneurons, such that SST interneuron silencing may partially disinhibit PV interneurons and vice versa ^41, 47, 48^. Some of these effects observed during optogenetic silencing of PV and SST interneurons may therefore reflect indirect circuit effects in addition to the direct loss of the targeted interneuron subtype. Future experiments employing simultaneous imaging of PV and SST interneuron activity during optogenetic perturbations will be needed to fully dissect these reciprocal inhibitory interactions. Relatedly, SST interneuron silencing paradoxically improved overall navigation efficiency, reflected in significantly reduced trajectory distance and latency (Fig. 7e,g). While consistent with PV disinhibition following removal of SST-mediated inhibition, we cannot exclude alternative explanations including nonspecific effects on locomotion speed or motivational state, which were not directly assessed here and require further investigation. Finally, while the barrier-detour task revealed a behaviorally significant double dissociation between PV and SST interneuron silencing, the modest effect sizes observed suggest that this single-session paradigm may not fully capture the functional consequences of disrupting egocentric precision and stability. Tasks requiring extended navigation over multiple sessions, or paradigms that more directly tax egocentric integration over longer timescales, may reveal stronger and more dissociable behavioral phenotypes in future studies.

Together, these findings reveal a division of labor between inhibitory microcircuits that allows RSC to integrate self-motion signals while maintaining stable egocentric spatial representations. PV interneurons enable rapid updating and precise alignment of egocentric representations with ongoing movement, whereas SST interneurons stabilize these representations over longer timescales and anchor them to environmental structure. Such dissociable inhibitory mechanisms may represent a general circuit strategy through which cortical networks balance flexibility and stability during dynamic spatial navigation.

## Acknowledgement

This work was supported by the National Research Foundation of Korea (NRF) grant RS-2024-00341894 (J.K.), the Institute of Information & Communications Technology Planning & Evaluation (IITP)-ITRC (information Technology Research Center) grant funded by the Korea government (MSIT) IITP-2025-RS-2024-00436857 (J.K.) and by the New Faculty Startup Fund from Seoul National University (J.K.).

## Author contributions

J.K. conceptualized the study and supervised the project. D.O. and J.Y. performed surgeries and experiments for calcium imaging. D.O. performed data analyses and generated all figures. J.Y. conducted initial analysis egocentric spatial representations in PV interneurons. J.S. performed optic fiber implant surgeries. D.O. and J.S. designed and analyzed the barrier-detour behavioral experiments and performed histological verification. J.K. wrote the manuscript. All authors reviewed and approved the manuscript. D.O. and J.Y. contributed equally to this work.

## References

1. Keshavarzi, S., et al. Multisensory coding of angular head velocity in the retrosplenial cortex. Neuron 110, 532–543 e539 (2022).

2. Taube, J.S. The head direction signal: origins and sensory-motor integration. Annu Rev Neurosci 30, 181–207 (2007).

3. O’Keefe, J. & Dostrovsky, J. The hippocampus as a spatial map. Preliminary evidence from unit activity in the freely-moving rat. Brain Res 34, 171–175 (1971).

4. Hafting, T., Fyhn, M., Molden, S., Moser, M.B. & Moser, E.I. Microstructure of a spatial map in the entorhinal cortex. Nature 436, 801–806 (2005).

5. Vann, S.D., Aggleton, J.P. & Maguire, E.A. What does the retrosplenial cortex do? Nat Rev Neurosci 10, 792–802 (2009).

6. Byrne, P., Becker, S. & Burgess, N. Remembering the past and imagining the future: a neural model of spatial memory and imagery. Psychol Rev 114, 340–375 (2007).

7. Vann, S.D. & Aggleton, J.P. Extensive cytotoxic lesions of the rat retrosplenial cortex reveal consistent deficits on tasks that tax allocentric spatial memory. Behav Neurosci 116, 85–94 (2002).

8. Maguire, E.A. The retrosplenial contribution to human navigation: a review of lesion and neuroimaging findings. Scand J Psychol 42, 225–238 (2001).

9. Clark, B.J., Bassett, J.P., Wang, S.S. & Taube, J.S. Impaired head direction cell representation in the anterodorsal thalamus after lesions of the retrosplenial cortex. J Neurosci 30, 5289–5302 (2010).

10. Alexander, A.S., Place, R., Starrett, M.J., Chrastil, E.R. & Nitz, D.A. Rethinking retrosplenial cortex: Perspectives and predictions. Neuron 111, 150–175 (2023).

11. Epstein, R.A., Patai, E.Z., Julian, J.B. & Spiers, H.J. The cognitive map in humans: spatial navigation and beyond. Nat Neurosci 20, 1504–1513 (2017).

12. Miller, A.M., Vedder, L.C., Law, L.M. & Smith, D.M. Cues, context, and long-term memory: the role of the retrosplenial cortex in spatial cognition. Front Hum Neurosci 8, 586 (2014).

13. Alexander, A.S. & Nitz, D.A. Retrosplenial cortex maps the conjunction of internal and external spaces. Nat Neurosci 18, 1143–1151 (2015).

14. Alexander, A.S., et al. Egocentric boundary vector tuning of the retrosplenial cortex. Sci Adv 6, eaaz2322 (2020).

15. Park, K., Yeo, Y., Shin, K. & Kwag, J. Egocentric neural representation of geometric vertex in the retrosplenial cortex. Nat Commun 15, 7156 (2024).

16. Cheng, N., et al. Egocentric processing of items in spines, dendrites, and somas in the retrosplenial cortex. Neuron 112, 646–660 e648 (2024).

17. Fischer, L.F., Mojica Soto-Albors, R., Buck, F. & Harnett, M.T. Representation of visual landmarks in retrosplenial cortex. Elife 9 (2020).

18. LaChance, P.A. & Hasselmo, M.E. Distinct codes for environment structure and symmetry in postrhinal and retrosplenial cortices. Nat Commun 15, 8025 (2024).

19. Miller, A.M.P., Mau, W. & Smith, D.M. Retrosplenial Cortical Representations of Space and Future Goal Locations Develop with Learning. Curr Biol 29, 2083–2090 e2084 (2019).

20. McNaughton, B.L., Battaglia, F.P., Jensen, O., Moser, E.I. & Moser, M.B. Path integration and the neural basis of the ’cognitive map’. Nat Rev Neurosci 7, 663–678 (2006).

21. Cardin, J.A., et al. Driving fast-spiking cells induces gamma rhythm and controls sensory responses. Nature 459, 663–667 (2009).

22. Hu, H., Gan, J. & Jonas, P. Interneurons. Fast-spiking, parvalbumin(+) GABAergic interneurons: from cellular design to microcircuit function. Science 345, 1255263 (2014).

23. Sohal, V.S., Zhang, F., Yizhar, O. & Deisseroth, K. Parvalbumin neurons and gamma rhythms enhance cortical circuit performance. Nature 459, 698–702 (2009).

24. Kepecs, A. & Fishell, G. Interneuron cell types are fit to function. Nature 505, 318–326 (2014).

25. Royer, S., et al. Control of timing, rate and bursts of hippocampal place cells by dendritic and somatic inhibition. Nat Neurosci 15, 769–775 (2012).

26. Miao, C., Cao, Q., Moser, M.B. & Moser, E.I. Parvalbumin and Somatostatin Interneurons Control Different Space-Coding Networks in the Medial Entorhinal Cortex. Cell 171, 507–521 e517 (2017).

27. Buetfering, C., Allen, K. & Monyer, H. Parvalbumin interneurons provide grid cell-driven recurrent inhibition in the medial entorhinal cortex. Nat Neurosci 17, 710–718 (2014).

28. Lovett-Barron, M., et al. Regulation of neuronal input transformations by tunable dendritic inhibition. Nat Neurosci 15, 423–430, S421-423 (2012).

29. Urban-Ciecko, J. & Barth, A.L. Somatostatin-expressing neurons in cortical networks. Nat Rev Neurosci 17, 401–409 (2016).

30. Fuchs, E.C., et al. Local and Distant Input Controlling Excitation in Layer II of the Medial Entorhinal Cortex. Neuron 89, 194–208 (2016).

31. Cowansage, K.K., et al. Direct reactivation of a coherent neocortical memory of context. Neuron 84, 432–441 (2014).

32. Yamawaki, N., et al. Long-range inhibitory intersection of a retrosplenial thalamocortical circuit by apical tuft-targeting CA1 neurons. Nat Neurosci 22, 618–626 (2019).

33. Park, K., Kohl, M.M. & Kwag, J. Memory encoding and retrieval by retrosplenial parvalbumin interneurons are impaired in Alzheimer’s disease model mice. Curr Biol 34, 434–443 e434 (2024).

34. Hinman, J.R., Chapman, G.W. & Hasselmo, M.E. Neuronal representation of environmental boundaries in egocentric coordinates. Nat Commun 10, 2772 (2019).

35. Vedder, L.C., Miller, A.M.P., Harrison, M.B. & Smith, D.M. Retrosplenial Cortical Neurons Encode Navigational Cues, Trajectories and Reward Locations During Goal Directed Navigation. Cereb Cortex 27, 3713–3723 (2017).

36. Taube, J.S. Head direction cells recorded in the anterior thalamic nuclei of freely moving rats. J Neurosci 15, 70–86 (1995).

37. Stackman, R.W. & Taube, J.S. Firing properties of head direction cells in the rat anterior thalamic nucleus: dependence on vestibular input. J Neurosci 17, 4349–4358 (1997).

38. Shinder, M.E. & Taube, J.S. Active and passive movement are encoded equally by head direction cells in the anterodorsal thalamus. J Neurophysiol 106, 788–800 (2011).

39. Onorato, I., et al. Distinct roles of PV and Sst interneurons in visually induced gamma oscillations. Cell Rep 44, 115385 (2025).

40. Cardin, J.A. Inhibitory Interneurons Regulate Temporal Precision and Correlations in Cortical Circuits. Trends Neurosci 41, 689–700 (2018).

41. Jang, H.J., et al. Distinct roles of parvalbumin and somatostatin interneurons in gating the synchronization of spike times in the neocortex. Sci Adv 6, eaay5333 (2020).

42. Udakis, M., Claydon, M.D.B., Zhu, H.W., Oakes, E.C. & Mellor, J.R. Hippocampal OLM interneurons regulate CA1 place cell plasticity and remapping. Nat Commun 16, 9912 (2025).

43. Jazayeri, M. & Ostojic, S. Interpreting neural computations by examining intrinsic and embedding dimensionality of neural activity. Curr Opin Neurobiol 70, 113–120 (2021).

44. Burak, Y. & Fiete, I.R. Accurate path integration in continuous attractor network models of grid cells. PLoS Comput Biol 5, e1000291 (2009).

45. Hardcastle, K., Ganguli, S. & Giocomo, L.M. Environmental boundaries as an error correction mechanism for grid cells. Neuron 86, 827–839 (2015).

46. Samsonovich, A. & McNaughton, B.L. Path integration and cognitive mapping in a continuous attractor neural network model. J Neurosci 17, 5900–5920 (1997).

47. Pfeffer, C.K., Xue, M., He, M., Huang, Z.J. & Scanziani, M. Inhibition of inhibition in visual cortex: the logic of connections between molecularly distinct interneurons. Nat Neurosci 16, 1068–1076 (2013).

48. Cottam, J.C., Smith, S.L. & Hausser, M. Target-specific effects of somatostatin-expressing interneurons on neocortical visual processing. J Neurosci 33, 19567–19578 (2013).

