## Supplementary Materials for "Retrosplenial PV and SST interneurons shape egocentric spatial precision and stability"

**This file includes:**

Materials and Methods
References (1 – 23)
Extended Data Fig. 1 - 7

### Materials and Methods

**Animals**

Male PV-Cre (#017320, Jackson Laboratory, USA) and SST-Cre mice (#013044, Jackson Laboratory, USA) aged 4 - 9 months were used for all experiments. Mice were housed individually in a temperature- and humidity-controlled environment on a 12 h light/dark cycle with food and water *ad libitum*. All experiments were conducted in accordance with and under the approval of the Institutional Animal Care and Use Committee of Seoul National University (IACUC; SNU-230410-1-2 and SNU-240426-3-3).

**Stereotaxic viral injection and surgical implantation**

Mice were anesthetized with 2% isoflurane and secured in a stereotaxic frame (51730D, Stoelting, USA). Viral vectors were injected into the retrosplenial cortex (RSC) at three sites (AP: −2.5 mm, ML: −0.3 mm; DV: −0.8, −0.6, and −0.4 mm) using a stereotaxic injector (53311, Stoelting, USA). All viral vectors were diluted 1:1 in saline and delivered at a rate of 50 nl/min (total volume: 500 nl per site).

For cell-type–specific imaging of PV or SST interneurons, PV-Cre or SST-Cre mice received injections of AAV1-EF1a-DIO-GCaMP6s-P2A-nls-dTomato (#51082, Addgene, USA). For simultaneous $Ca^{2+}$ imaging of excitatory neurons and optogenetic silencing of PV or SST interneurons, AAV9-CaMKII-GCaMP6s-WPRE-SV40 (#107790, Addgene, USA) was co-injected with pAAV5-EF1a-DIO-eNpHR3.0-EYFP (#26966, Addgene, USA) into PV-Cre or SST-Cre mice.

For $Ca^{2+}$ imaging experiments, a craniotomy was performed above the RSC (AP: −2.5 mm, ML: ±0.3 mm), and a GRIN lens (1 mm diameter, 4 mm length; #1050-004605, Inscopix, USA) was slowly lowered to DV: −0.5 mm and secured with dental adhesive resin cement (Super-

Bond, Sun Medical, Japan). After 2–4 weeks of recovery, a magnetic baseplate (#1050-004638, Inscopix, USA) was positioned above the lens to obtain an optimal field of view (FOV) using a miniaturized microscope (nVoke 2.0, Inscopix, USA). Once the imaging plane was identified, the baseplate was permanently fixed, and mice were allowed to recover for at least one week before the start of imaging experiments.

For optogenetic silencing experiments, pAAV5-EF1a-DIO-eNpHR3.0-EYFP (#26966, Addgene, USA) was injected bilaterally into the RSC (AP: −2.5 mm; ML: ±0.3 mm; DV: −0.8, −0.6, −0.4, and −0.2 mm). Optical fibers (#R-FOC-L200C-50NA, RWD Life Science, China) were implanted bilaterally with fiber tips positioned at AP: −2.5 mm, ML: ±0.3 mm, DV: −0.4 mm. To accommodate the thickness of the fiber ferrule and avoid interference with the skull, fibers were implanted at an angle of ~10° relative to the vertical axis angled laterally away from the midline and secured with dental adhesive resin cement.

**In vivo $Ca^{2+}$ imaging and data preprocessing**

To perform cell-type-specific one-photon $Ca^{2+}$ imaging, GCaMP6s-expressing neurons were acquired using the nVoke 2.0 and Inscopix Data Acquisition software (nVoke, Inscopix, USA) at a sampling rate of 20 Hz (exposure time: 50 ms) while mice freely explored the square open-field chamber for 30 min. For each mouse, LED power (0.1–0.3 mW/mm$^2$), digital focus, gain, and field of view were adjusted to optimize signal quality based on GCaMP6s expression within the FOV, and the same parameters were maintained across sessions to enable longitudinal recordings from the same neuronal population. Raw $Ca^{2+}$ imaging data were preprocessed using Inscopix Data Processing Software (IDPS, Inscopix, USA) as described in previous studies [1,2]. Images were spatially downsampled by a factor of 2 and band-pass filtered (low cutoff: 0.005 Hz; high cutoff: 0.5 Hz). Motion correction was applied to all data using cell registration-based image registration method [2,3]. Fluorescence signals were expressed as $\Delta F/F_0$, where $\Delta F = F - F_0$, F is the fluorescence signal at each time point, and $F_0$ represents the baseline fluorescence level signal. Putative neurons and their $Ca^{2+}$ transients were identified using an extraction procedure for individual neuronal signals based on principal component analysis followed by independent component analysis (PCA–ICA) [1, 4]. Spatial components exhibiting neuronal morphology and temporally consistent $Ca^{2+}$ dynamics were retained for further analysis, whereas non-neuronal components were excluded. Putative spike events were inferred from denoised ΔF/F traces using the Online Active Set method to Infer Spikes (OASIS) [2, 5, 6], a deconvolution-based approach for estimating spike timing from calcium signals.

**Optogenetic silencing of PV and SST interneurons during $Ca^{2+}$ imaging of RSC excitatory neurons**

Simultaneous $Ca^{2+}$ imaging and optogenetic silencing were performed using a miniature integrated microscope system (nVoke 2.0, Inscopix, USA). Imaging parameters were kept identical to those used in control sessions. Optogenetic silencing was achieved using the built-in optogenetic LED in nVoke 2.0 (620 ± 30 nm, 5 $mW/mm^2$ at the imaging plane). Light was delivered in a 2-s on / 1-s off cycle during the entire designated silencing periods to silence eNpHR3.0-expressing PV or SST interneurons. This intermittent illumination protocol was used to reduce photobleaching, limit tissue heating, and prevent opsin desensitization that could occur during sustained illumination in one-photon $Ca^{2+}$ imaging experiments in freely behaving mice. Effective silencing of PV or SST interneurons was confirmed by changes in RSC excitatory neuron firing rates during light-on epochs relative to light-off epochs within the same session (Fig. 4c).

**Open field behavior experiments**

To perform one-photon $Ca^{2+}$ imaging during spatial exploration, mice were habituated to the recording setup and behavioral environment prior to imaging experiments. Mice underwent at least 3 days of handling before $Ca^{2+}$ imaging to allow them to adapt to the head-mounted miniaturized microscope and the commutator system (Inscopix, USA). Fixed illumination (23 lux) was maintained in a sound-attenuated recording room to minimize variability in sensory conditions across experimental sessions. Experiments were performed in a custom-built square open-field chamber (35 × 35 cm), in which small food crumbs (Froot Loops, Kellogg's, USA) were scattered to promote uniform spatial exploration of the arena. Each open field session lasted 30 min. Behavioral data, including the position and head direction of each mouse, were acquired using EthoVision XT 16 software (Noldus, Netherlands) at a sampling rate of 15 Hz. Head direction and body position tracking were manually verified to ensure accurate detection.

**Goal-directed navigation task experiment**

To investigate the role of PV and SST interneurons in goal-directed navigation, a goal-directed navigation task was adapted based on a previously reported barrier detour paradigm [7]. Mice were food-restricted and maintained at 80 – 85% of their free-feeding body weight throughout the task period. The goal-directed navigation task was conducted in a square open chamber (50 × 50 cm, wall height 30 cm). A 15 × 15 cm start box was mounted at the midpoint of one wall and connected to the arena by a 5-cm-wide opening. Access from the start box to the chamber was controlled by a sliding door, which remained closed during the inter-trial period and was opened at trial onset to allow the animal to enter the chamber. The reward (Froot Loops, Kellogg's, USA) location was positioned at the midpoint of the wall opposite the start box. An opaque barrier (40 cm long, 0.5 cm thick) extending from one wall and leaving a 10 cm gap on the opposite side, was positioned in the center of the arena to create a partial barrier between the start and reward locations.

Goal-directed navigation task training was performed over 4 consecutive days, up to 25 daily trials, with the number of trials varying across mice (13 – 25 trials per mouse per day). At the beginning of each trial, the mouse was placed into the start box and released by opening the sliding door. A trial ended when the mouse reached and consumed the reward. Following reward acquisition, the mouse was guided back to the start box, where a second reward was delivered in the start box.

On the fifth day, PV or SST interneurons were optogenetically silenced bilaterally throughout each trial by delivering 595 nm light stimulation (5 $mW/mm^2$ at tip) using a fiber-coupled LED (M595F2, Thorlabs, USA) connected to the implanted optic fiber cannula (#R-FOC-L200C-50NA, RWD Life Science, China) during the barrier-detour, goal-directed navigation task.

**Firing rate maps and spatial stability analysis**

To quantify the spatial receptive fields and spatial stability of RSC neurons during free exploration, $Ca^{2+}$ imaging data and behavioral tracking data obtained with EthoVision XT 16 (Noldus, Netherlands) were synchronized using a Noldus IO Box (Mini USB-IO Box, Noldus, Netherlands) and analyzed using custom-written MATLAB code (MathWorks, USA). To characterize the spatial receptive fields of GCaMP6s-expressing RSC neurons, the position of each mouse in the square open chamber was discretized into spatial bins of 1.5 cm × 1.5 cm. For each spatial bin, neuronal firing rate was computed by dividing the total number of deconvolved spike events by the total occupancy time in that bin [8, 9]. Spatial bins with occupancy times shorter than 67 ms (corresponding to the temporal resolution of the behavioral tracking data) and spike events occurring when the animal's instantaneous velocity was below 2 cm/s were excluded from analysis.

The resulting firing rate maps were smoothed using a boxcar averaging filter (5 × 5 bins) followed by a Gaussian kernel ($\sigma = 1$ bin).[10, 11] Spatial receptive fields were defined as contiguous regions of at least 20 $cm^2$ in which the firing rate exceeded 30% of the peak firing rate of the smoothed firing rate map, adapted from previously reported criteria for spatially tuned neurons [10, 11].

Spatial stability of spatial receptive field was assessed by calculating the Pearson correlation coefficient between firing rate maps constructed from the first half (0 – 15 min) and the second half (15 – 30 min) of each recording session [8, 10, 12]. To determine statistical significance, a shuffled distribution of correlation coefficients was generated by circularly time-shifting each neuron's spike train by a random offset relative to the animal's trajectory, repeated 500 times. This procedure preserved the temporal structure of both $Ca^{2+}$ signals and behavioral data while disrupting their temporal alignment. Neurons with spatial correlation coefficients exceeding the 99th percentile of the shuffled distribution were classified as spatially stable.

**Egocentric boundary cell analysis**

To identify egocentric boundary cells, an analytical framework previously established for egocentric boundary coding in the RSC was adopted [8, 12-14]. At each behavioral time point, the egocentric bearing ($\theta_{\mathrm{ego}}$) and egocentric distance ($D_{\mathrm{ego}}$) to the nearest environmental boundary were calculated relative to the animal's instantaneous head direction and position.

Behavioral samples were binned in a two-dimensional egocentric polar coordinate system consisting of angular bins of 15° and radial distance bins of 1.5 cm. For each neuron, the firing rate was computed as the number of spike events divided by the occupancy duration in each bin. Only bins with occupancy of at least one behavioral frame (≥ 66 ms) were included in the analysis. Egocentric firing rate maps were smoothed using a two-dimensional Gaussian kernel ($\sigma = 0.7$ pixels). To quantify egocentric boundary tuning, the mean resultant egocentric boundary vector was computed as follows [8, 12, 14]:

$$\text{Egocentric } boundary - vector = \left(\sum_{\theta=1}^{n}\sum_{D=1}^{m} F_{\theta,D} * e^{i\theta}\right)/(n * m * \bar{F})$$

where $\theta$ denotes egocentric bearing relative to the boundary, $D$ denotes egocentric distance to the boundary, and $F_{\theta,D}$ represents the firing rate in each angular–distance bin. $n$ and $m$ indicate the total number of angular and distance bins, respectively. $e^{i\theta}$ is the complex unit vector at angle $\theta$, and $\bar{F}$ is the mean firing rate across all bins. The magnitude of this complex vector was defined as the egocentric boundary vector length (VL), providing a scalar measure of egocentric tuning strength ranging from 0 (no tuning) to 1 (perfect tuning).

The preferred egocentric bearing was defined as the mean resultant angle (MRA):

$$\text{MRA} = \text{arctan2}(\text{imag}(\text{egocentric boundary vector}))/\text{real}(\text{egocentric boundary vector}))$$

The preferred egocentric distance was estimated by extracting the radial firing rate profile along the preferred egocentric bearing and fitting it with a Weibull function [8, 12]. The distance corresponding to the peak of the fitted function was defined as the neuron's preferred $D_{\text{ego}}$.

Statistical significance was assessed using a shuffle procedure in which each neuron's spike train was circularly time-shifted relative to the animal's behavioral trajectory by a random temporal offset, repeated 500 times. For each shuffle, the egocentric firing rate map and vector length were recomputed. Neurons were classified as egocentric boundary cells if their observed vector length exceeded the 99$^{\text{th}}$ percentile of the shuffled distribution.

**Border score analysis**

To quantify the extent to which neuronal firing was anchored to environmental boundaries, border cell analysis was performed using an established framework previously applied to spatial boundary coding in the RSC [10]. For each neuron, spatial firing-rate maps were constructed by binning the animal's position into two-dimensional spatial grids and calculating the firing rate within each bin. The resulting maps were smoothed using a Gaussian kernel ($\sigma = 1$) to reduce sampling noise and improve spatial receptive field estimation. Putative border fields were identified as contiguous regions of elevated firing that directly contacted an environmental wall and exceeded 30% of the neuron's peak firing rate in the smoothed rate map. For each wall of the enclosure, border coverage was quantified as the proportion of spatial bins along that wall that were occupied by the identified border field. The maximum coverage value across the four walls was defined as the border coverage score ($c$). The mean firing distance ($d$) was calculated by measuring the mean distance from the spatial bins within the border field to the nearest environmental boundary, weighted by the normalized firing rate of each bin. To allow comparison across environments of different sizes, the weighted distance was normalized by the maximum possible distance to the nearest border, yielding values between 0 and 1. The border score ($b$) was then calculated by comparing $c$ and $d$, following a method adapted from previous studies:

$$b = (c - d)/(c + d)$$

where values near +1 indicate firing fields tightly aligned with an environmental border, and values near −1 indicate fields centered away from all borders. Statistical significance was assessed via a shuffle procedure in which spike trains were circularly time-shifted relative to the animal's trajectory to generate a null distribution of border scores. Neurons whose observed border score exceeded the 99$^{th}$ percentile of the shuffle distribution were classified as border cells.

**Self-motion tuning analysis**

To identify neurons tuned to self-motion variables, tuning to angular head velocity (AHV) and linear locomotion speed was quantified using open-field recordings, based on the moment-to-moment relationship between each behavioral variable and neuronal firing rate, following previously established procedures [15-17]. Head direction was first unwrapped and smoothed using a Gaussian kernel ($\sigma = 200$ ms) to reduce tracking noise. AHV was then computed as the temporal derivative of the smoothed head-direction signal at each time point using a 200-ms sliding window. Positive and negative AHV values corresponded to rightward and leftward head turns, respectively. For each neuron, AHV tuning curves were constructed by binning AHV into 6°/s bins, spanning $|AHV| \leq 300°/s$. The firing rate in each bin was calculated as the number of spikes divided by the occupancy time. AHV tuning strength was quantified using the Pearson correlation coefficient between firing rate and AHV across all bins [17]. For directional analyses, correlations were computed separately for right-turn and left-head turns, and the absolute correlation value was used as the AHV score.

Linear locomotion speed was computed from the animal's two-dimensional position as the displacement between consecutive video frames divided by the sampling interval, and the resulting speed trace was smoothed using a sliding mean filter. Speed tuning curves were constructed by binning locomotion speed at 1 cm/s bins within the range of 0–20 cm/s, and firing rate in each bin was calculated as the spike count divided by the occupancy time. Speed scores were defined as the absolute Pearson correlation coefficient between firing rate and speed across bins [15, 16].

To assess statistical significance, shuffle distributions were generated independently for each neuron by circularly time-shifting spike trains relative to the behavioral time series and computing correlation values for each shuffle (500 iterations). Neurons were classified as AHV-tuned or speed-tuned if the observed correlation exceeded the 99th percentile of the shuffled distribution.

To characterize self-motion tuning gradients, linear regression was performed between firing rate and AHV separately for right-turn and left-turn movements, and between firing rate and speed within the sampled speed range. Behavioral samples with extreme values (>300°/s for AHV, >20 cm/s for speed) were excluded due to low sampling probability.

**Population vector distance analysis**

To quantify population-level similarity across egocentric bearing conditions, population vector analysis was performed [18, 19]. Neuronal activity was Gaussian-smoothed ($\sigma = 200$ ms), and egocentric bearings were discretized into 15° bins. For each bearing bin, population vectors were constructed by averaging neuronal activity across all time points assigned to that bin. Population vectors corresponding to different bearing bins were then compared using cosine distance. To align angular bins to a common egocentric reference frame, the resulting distance matrix was circularly shifted such that the bin corresponding to 0° egocentric bearing was centered, yielding an egocentric bearing-aligned population distance matrix. To quantify how population similarity varied as a function of egocentric bearing difference, cosine distance values were compared between neurons with similar egocentric bearing preferences (0° difference) and neurons with opposing preferences (180° difference).

**Synchrony analysis**

Neuronal synchrony was assessed using pairwise correlation analysis of deconvolved $Ca^{2+}$ spike trains obtained during 30-min open-field exploration. For each neuron, the deconvolved spike train was smoothed by convolution with a Gaussian kernel (standard deviation = 200 ms) to generate a continuous activity trace. All smoothed traces were z-scored within each session to account for differences in baseline activity and variance across neurons. Pairwise synchrony was quantified using the Pearson correlation coefficient computed between their smoothed activity traces of all neuron pairs across the entire recording session.

**Temporal lag analysis**

To characterize the temporal relationship between interneuronal population activity and self-motion variables, time-lagged correlation analysis was performed following established methods [14]. Spike trains were first binned at the behavioral sampling rate (15 Hz; 67 ms per bin) and smoothed using a Gaussian kernel (standard deviation = 200 ms) to obtain continuous firing rate estimates. AHV and running speed traces were then circularly shifted relative to the population neuronal activity vectors over a temporal window ranging from −2 to +2 s. At each temporal lag, the Pearson correlation coefficient between neuronal activity and the behavioral variable was computed. For AHV, correlations were calculated separately for positive (right turn) and negative (left turn) velocity epochs to account for directional asymmetry, and the larger correlation coefficients was taken as the coupling strength at each lag. For speed, correlations were computed using the full speed trace. The lag–correlation function was used to determine the temporal alignment between population neuronal activity and self-motion variables. The temporal lag corresponding to the peak correlation coefficient was defined as the preferred coupling latency, indicating whether neuronal activity preceded, coincided with, or followed the self-motion signal.

**Population clustering analysis**

To examine whether neuronal population activity exhibits egocentric bearing-dependent clustering structure as a function of differences in preferred egocentric bearing, neuronal activity was Gaussian-smoothed ($\sigma$ = 200 ms). To ensure stable behavioral sampling and minimize potential non-stationarity across long recordings, analyses were restricted to a continuous 10-min segment of each session with sufficient behavioral coverage of egocentric bearings. Principal component analysis (PCA) was then performed on the neuronal activity matrix (time bins × neurons), and the top ten principal components were retained to capture the dominant population structure while reducing dimensionality. Pairwise Pearson correlation coefficients between all neuron pairs were then computed in PC-reduced space as a measure of functional similarity. For each neuron pair, the absolute difference in preferred egocentric bearing was calculated, and neuron pairs were grouped into five angular bins spanning 0° to 180° to assess clustering as a function of egocentric tuning similarity.

**Low-dimensional embedding of neural activity**

To visualize the low-dimensional structure of neural population activity, a two-step dimensionality reduction approach was implemented based on a previously published method [20]. Binarized, deconvolved spike activity from all neurons in each session (time bin = 67 ms) was convolved with a Gaussian kernel ($\sigma$ = 200 ms) to obtain smoothed firing rate estimates. The smoothed activity of each neuron was then z-scored across time. Dimensionality reduction was subsequently performed on the resulting data matrix (time bins × neurons). Principal component analysis (PCA) was first applied to reduce noise and capture dominant variance across the neural population. Principal components explaining 70% of the total variance were selected and used as input to Uniform Manifold Approximation and Projection (UMAP), which generated a low-dimensional three-dimensional embedding. UMAP parameters were set to min_dist = 0.1, n_neighbors = 50, and n_components = 3. These parameters were held constant across all session and conditions to ensure comparability.

**Bayesian decoding of egocentric bearing**

To assess whether egocentric bearing relative to the nearest boundary could be reconstructed from neuronal population activity, a Bayesian decoder was applied to deconvolved $Ca^{2+}$ signals[21]. Egocentric bearing was discretized into angular bins of 15°, consistent with the binning used for egocentric firing rate map construction. Decoding performance was evaluated using 10-fold cross-validation, in which data were partitioned into training and test sets, and the decoder was trained on the training set and evaluated on the held-out test set. Neural activity was aligned to behavioral time points, and at each time point, the posterior probability of each angular bin was computed based on the observed population activity using Bayes' theorem. The angular bin with the highest posterior probability was selected as the decoded egocentric bearing. Decoding accuracy was quantified by calculating the mean absolute difference between the decoded and true egocentric bearing across all time points in the test set, representing the mean angular decoding error.

**Barrier-detour goal-directed navigation analysis**

To evaluate goal-directed learning and determine whether optogenetic silencing of PV and SST interneurons alters goal-directed navigation behavior, the animal's nose and body center position were extracted using DeepLabCut from video recordings acquired at 7.5 Hz [22]. The start of each trial was defined as the moment the animal's nose crossed the opening of the sliding door that connected the start box (located at x = 0 cm) to the open square chamber where the reward (located at x = 50 cm) was located. Reward arrival was defined as the first moment the animal began consuming the reward. The time and distance traveled between these two events were calculated for each trial and averaged across trials per day to assess learning progression and navigation efficiency.

To quantify egocentric orientation during the approach to the barrier gap end, $\theta_{\mathrm{ego}}$ was computed across x = 0–20 cm, beginning at trial onset and ending 5 cm before the barrier (located at x = 25 cm) to exclude trajectory adjustments during barrier passage. This range was divided into twenty equally spaced spatial bins (bin width = 1 cm). $\theta_{\mathrm{ego\_initial}}$ was defined as the mean $\theta_{\mathrm{ego}}$ within the first spatial bin (x = 0–1 cm), reflecting egocentric orientation at navigation onset, and $\theta_{ego\_final}$ as the mean $\theta_{ego}$ within the last spatial bin (x = 19–20 cm), reflecting egocentric orientation immediately prior to barrier passage.

Analysis was restricted to the approach phase from start towards the barrier (x = 0–20 cm), because the post-barrier trajectory toward the reward is visually guided—the reward location is directly visible once the animal passes through the barrier gap—and therefore does not require egocentric spatial computation. Egocentric orientation analyses were thus confined to the non-visually-guided approach phase, where egocentric bearing toward the barrier gap end provides the primary navigational reference. Although homing trajectories—from the reward location back to the start box—were recorded, these were not analyzed quantitatively as animals exhibited highly variable return paths that precluded reliable egocentric bearing analysis, likely reflecting the availability of multiple viable return routes in the open arena.

**Histology**

To confirm the position of the GRIN lens track and optic fiber implant track, as well as the expression of GCaMP6s and eNpHR3.0 in RSC neurons, mice were first deeply anesthetized with Avertin (T48402, Sigma-Aldrich, USA) and transcardially perfused with 10 mL of ice-cold 4% paraformaldehyde (158127, Sigma-Aldrich, USA). Brains were carefully extracted and post-fixed overnight in 4% paraformaldehyde, and then transferred to a 30% sucrose solution in phosphate-buffered saline (PBS) for cryoprotection for 24–48 h. Brain tissues were embedded in optimum cutting temperature compound (Tissue-Tek O.C.T. Compound, Sakura Finetek, Japan) and frozen at −80 °C. Coronal sections (50 μm thick) were obtained using a cryostat (YD-2235, Jinhua Yidi Medical Appliance Co., China). Sections were washed three times with PBS to remove residual OCT compound and mounted on glass slides using antifade mounting medium containing DAPI (Vectashield, Vector Laboratories, USA). GRIN lens and optic fiber implant locations within RSC were identified based on the position of the GRIN lens track or the optical fiber cannula tip, respectively. Expression of GCaMP6s in excitatory neurons, PV interneurons, and SST interneurons or eNpHR3.0 in PV or SST interneurons was verified by detecting fluorescence signals using a confocal laser scanning microscope (LSM-700, Zeiss, Germany) and a fluorescence microscope (DM2500, Leica, Germany). Simultaneous confocal verification of GCaMP6s and eNpHR3.0-eYFP expression was not feasible in combined imaging-silencing experiments in Fig. 4 due to the overlapping emission spectra of GCaMP6s (~510 nm) and eYFP (~ 527 nm); expression of each fluorophore was therefore confirmed in separate tissue sections. Cell-type-specific eNpHR3.0-eYFP expression in PV and SST interneurons was confirmed histologically in Fig. 7b. GCaMP6s expression in RSC excitatory neurons was verified in a previously published study from our laboratory [23].

**Statistics**

Statistical significance was assessed using paired or unpaired Student's *t*-test or the nonparametric Wilcoxon rank-sum test or Watson-Williams test, as appropriate. A $p$ value less than 0.05 was considered statistically significant. Data are presented as mean ± SEM unless otherwise stated.

#### References


1. Resendez, S.L., *et al.* Visualization of cortical, subcortical and deep brain neural circuit dynamics during naturalistic mammalian behavior with head-mounted microscopes and chronically implanted lenses. *Nat Protoc* **11**, 566-597 (2016).
2. Stringer, C. & Pachitariu, M. Computational processing of neural recordings from calcium imaging data. *Curr Opin Neurobiol* **55**, 22-31 (2019).
3. Thevenaz, P., Ruttimann, U.E. & Unser, M. A pyramid approach to subpixel registration based on intensity. *IEEE Trans Image Process* **7**, 27-41 (1998).
4. Stamatakis, A.M., *et al.* Miniature microscopes for manipulating and recording in vivo brain activity. *Microscopy (Oxf)* **70**, 399-414 (2021).
5. Friedrich, J., Zhou, P. & Paninski, L. Fast online deconvolution of calcium imaging data. *PLoS Comput Biol* **13**, e1005423 (2017).
6. Pachitariu, M., Stringer, C. & Harris, K.D. Robustness of Spike Deconvolution for Neuronal Calcium Imaging. *J Neurosci* **38**, 7976-7985 (2018).
7. Shamash, P., *et al.* Mice learn multi-step routes by memorizing subgoal locations. *Nat Neurosci* **24**, 1270-1279 (2021).
8. Park, K., Yeo, Y., Shin, K. & Kwag, J. Egocentric neural representation of geometric vertex in the retrosplenial cortex. *Nat Commun* **15**, 7156 (2024).
9. Sun, Y., Nitz, D.A., Xu, X. & Giocomo, L.M. Subicular neurons encode concave and convex geometries. *Nature* **627**, 821-829 (2024).
10. Solstad, T., Boccara, C.N., Kropff, E., Moser, M.B. & Moser, E.I. Representation of geometric borders in the entorhinal cortex. *Science* **322**, 1865-1868 (2008).
11. Wills, T.J., Lever, C., Cacucci, F., Burgess, N. & O'Keefe, J. Attractor dynamics in the hippocampal representation of the local environment. *Science* **308**, 873-876 (2005).
12. Alexander, A.S., *et al.* Egocentric boundary vector tuning of the retrosplenial cortex. *Sci Adv* **6**, eaaz2322 (2020).
13. Hinman, J.R., Chapman, G.W. & Hasselmo, M.E. Neuronal representation of environmental boundaries in egocentric coordinates. *Nat Commun* **10**, 2772 (2019).
14. van Wijngaarden, J.B., Babl, S.S. & Ito, H.T. Entorhinal-retrosplenial circuits for allocentric-egocentric transformation of boundary coding. *Elife* **9** (2020).
15. Kropff, E., Carmichael, J.E., Moser, M.B. & Moser, E.I. Speed cells in the medial entorhinal cortex. *Nature* **523**, 419-424 (2015).
16. Campbell, M.G., *et al.* Principles governing the integration of landmark and self-motion cues in entorhinal cortical codes for navigation. *Nat Neurosci* **21**, 1096-1106 (2018).
17. Keshavarzi, S., *et al.* Multisensory coding of angular head velocity in the retrosplenial cortex. *Neuron* **110**, 532-543 e539 (2022).
18. Lee, I., Yoganarasimha, D., Rao, G. & Knierim, J.J. Comparison of population coherence of place cells in hippocampal subfields CA1 and CA3. *Nature* **430**, 456-459 (2004).
19. Pastalkova, E., Itskov, V., Amarasingham, A. & Buzsaki, G. Internally generated cell assembly sequences in the rat hippocampus. *Science* **321**, 1322-1327 (2008).
20. Gardner, R.J., *et al.* Toroidal topology of population activity in grid cells. *Nature* **602**, 123-128 (2022).
21. Brown, E.N., Frank, L.M., Tang, D., Quirk, M.C. & Wilson, M.A. A statistical paradigm for neural spike train decoding applied to position prediction from ensemble firing patterns of rat hippocampal place cells. *J Neurosci* **18**, 7411-7425 (1998).
22. Mathis, A., *et al.* DeepLabCut: markerless pose estimation of user-defined body parts with deep learning. *Nat Neurosci* **21**, 1281-1289 (2018).
23. Park, K., Kohl, M.M. & Kwag, J. Memory encoding and retrieval by retrosplenial parvalbumin interneurons are impaired in Alzheimer's disease model mice. *Curr Biol* **34**, 434-443 e434 (2024).

#### Extended Data Fig. 1

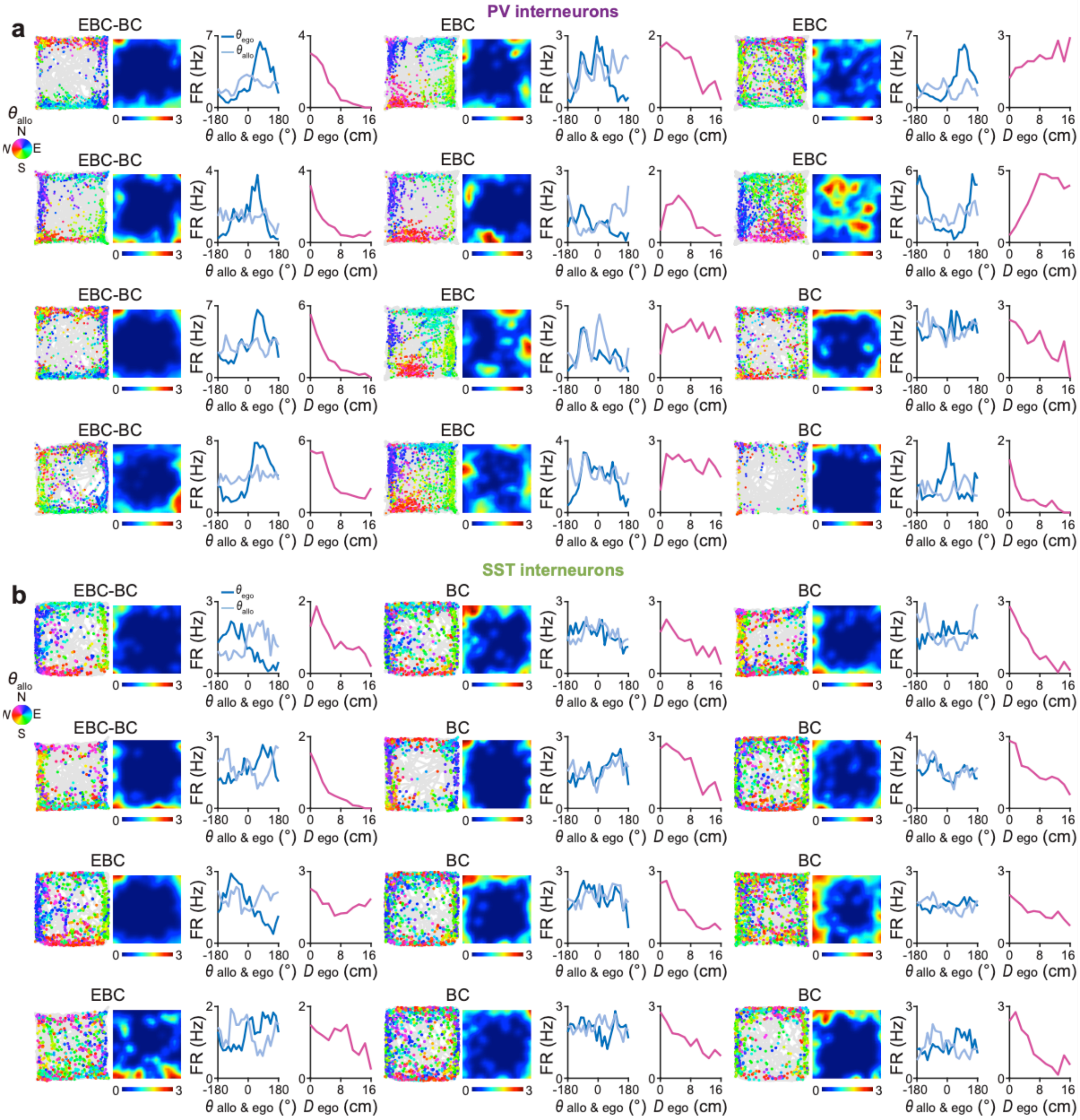


**Extended Data Fig. 1 Representative examples of egocentric boundary cells and border cells in PV and SST interneurons in the retrosplenial cortex.**

**a-b,** Representative examples of egocentric boundary cells (EBCs), border cells (BCs), and conjunctive EBC–BC neurons recorded from PV (**a**) and SST interneurons (**b**) during 30-min free exploration in a square open chamber. (Left) Spike-trajectory plots showing the animal's trajectory (gray) and spike locations (colored dots). Each spike is color-coded by allocentric head direction ($\theta_{allo}$) (inset, North: N; East: E; South: S; and West: W). (middle left) Corresponding spatial firing rate maps. (middle right) Angular tuning curves showing the preferred egocentric bearing relative to the closest boundary ($\theta_{ego}$, dark blue) and allocentric head direction $\theta_{allo}$ (light blue). (right) Egocentric distance tuning curves to the closest boundary ($D_{ego}$, magenta). n = 328 PV interneurons from eleven PV-Cre mice; n = 222 SST interneurons from six SST-Cre mice.

#### Extended Data Fig. 2

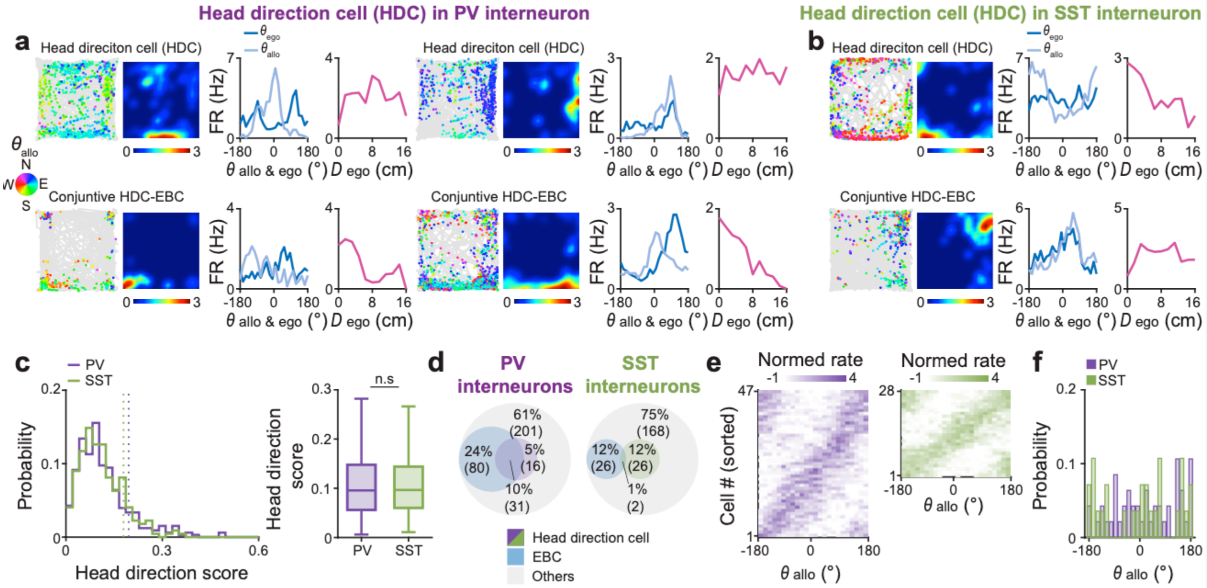


**Extended Data Fig. 2 Head direction tuning of PV and SST interneurons in the retrosplenial cortex**

**a-b,** Representative examples of head direction cells (HDCs), egocentric boundary cells (EBCs), and conjunctive HDC–EBC neurons recorded from PV (**a**) and SST (**b**) interneurons during 30-min free exploration in a square open chamber. (left) Spike-trajectory plots showing the animal's trajectory (gray) and spike locations (colored dots). Each spike is color-coded by allocentric head direction ($\theta_{allo}$) (inset, North: N; East: E; South: S; and West: W). (middle left) Corresponding spatial firing rate maps. (Middle right) Angular tuning curves showing the preferred egocentric bearing relative to the closest boundary ($\theta_{ego}$, dark blue) and allocentric head direction $\theta_{allo}$ (light blue). (right) Egocentric distance tuning curves to the closest boundary ($D_{ego}$. magenta).

**c,** (left) Probability distributions of head direction scores for PV (purple) and SST interneurons (green). Vertical dotted line: $99^{th}$ percentile of randomly shuffled shuffled head direction score of PV (purple) and SST interneurons (green). (right) Box plot of head direction score in PV (purple) and SST interneurons (green).

**d,** Venn diagrams showing the proportions of EBCs (blue), HDCs (PV interneurons: purple, SST interneurons: green), and other cell types (gray) in PV (left) and SST interneurons (right).

**e,** Normalized firing rate maps aligned to $\theta_{allo}$.

**f,** Probability distribution of preferred $\theta_{allo}$ in PV (purple) and SST interneurons (green).

Box plot (**c**) shows 25th (lower box line), 50th (middle line), and 75th (upper box line) percentile values; whiskers indicate minimum and maximum values. Two-sided Wilcoxon rank-sum test, $P = 0.7418$ (**c**). n = 328 PV interneurons from eleven PV-Cre mice; n = 222 SST interneurons from six SST-Cre mice.

Extended Data Fig. 3

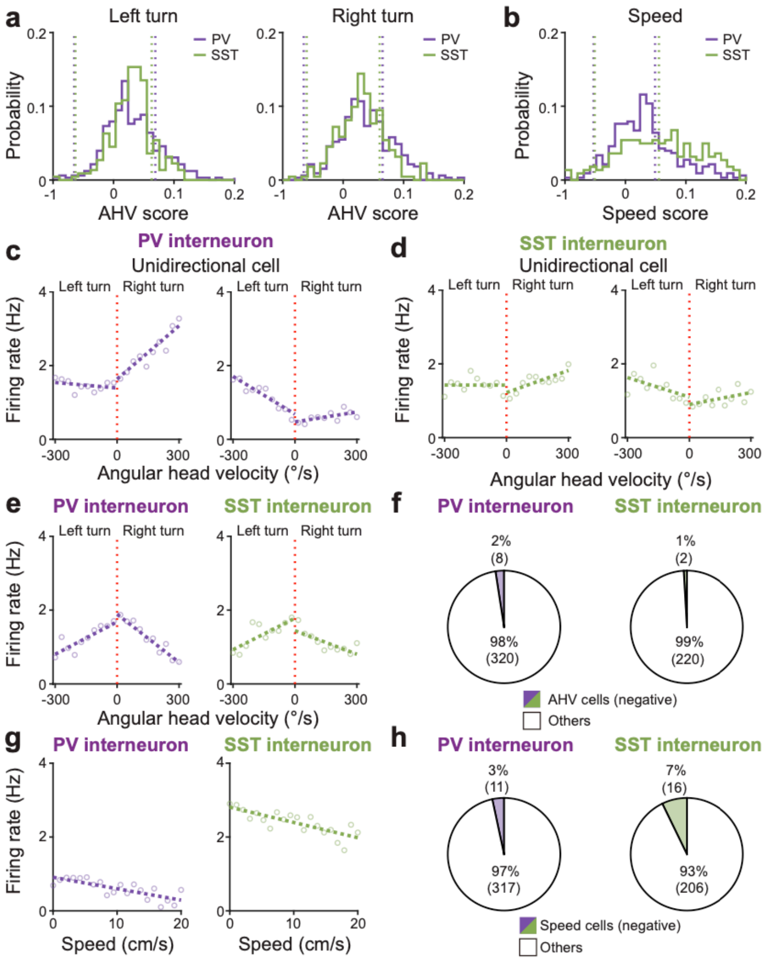


**Extended Data Fig. 3 Negatively tuned self-motion cells in PV and SST interneurons.** **a,** Probability distributions of angular head velocity (AHV) scores for left-turn and right-turn in PV (purple) and SST interneurons (green). Vertical red dotted line: 0°/s.

**b,** Same as **a,** but for locomotion speed scores.

**c-d,** Representative AHV tuning curves of negatively tuned AHV cells showing unidirectional suppression during left-turn and right-turn for PV (**c**) and SST interneurons (**d**) during left turns and right turns.

**e,** Population-averaged AHV tuning curves of negatively tuned AHV cells aligned to 0°/s for PV (purple) and SST interneurons (green).

**f,** Venn diagrams showing the proportions of negatively tuned AHV cells and other cell types in PV (left) and SST interneurons (right).

**g-h,** Same as **e-f,** but for negatively tuned locomotion speed cells.

n = 328 PV interneurons from eleven PV-Cre mice; n = 222 SST interneurons from six SST-Cre mice.

Extended Data Fig. 4

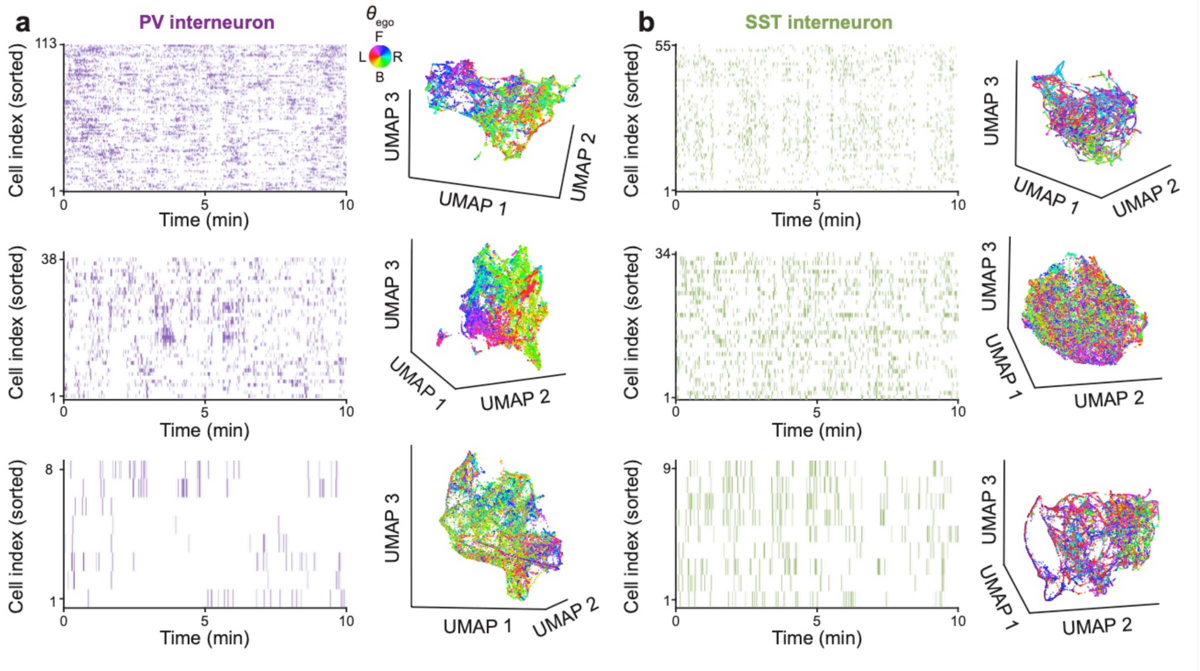


**Extended Data Fig. 4 Additional representative examples of low-dimensional population embeddings in PV and SST interneurons.**

**a,** (left) Example spike raster plots of PV interneurons during free exploration. (right) UMAP embedding of population activity for PV interneurons, colored by preferred egocentric bearing ($\theta_{ego}$).

**b,** Same as **a,** but for SST interneurons. n = 328 PV interneurons from eleven PV-Cre mice; n = 222 SST interneurons from six SST-Cre mice.

Extended Data Fig. 5

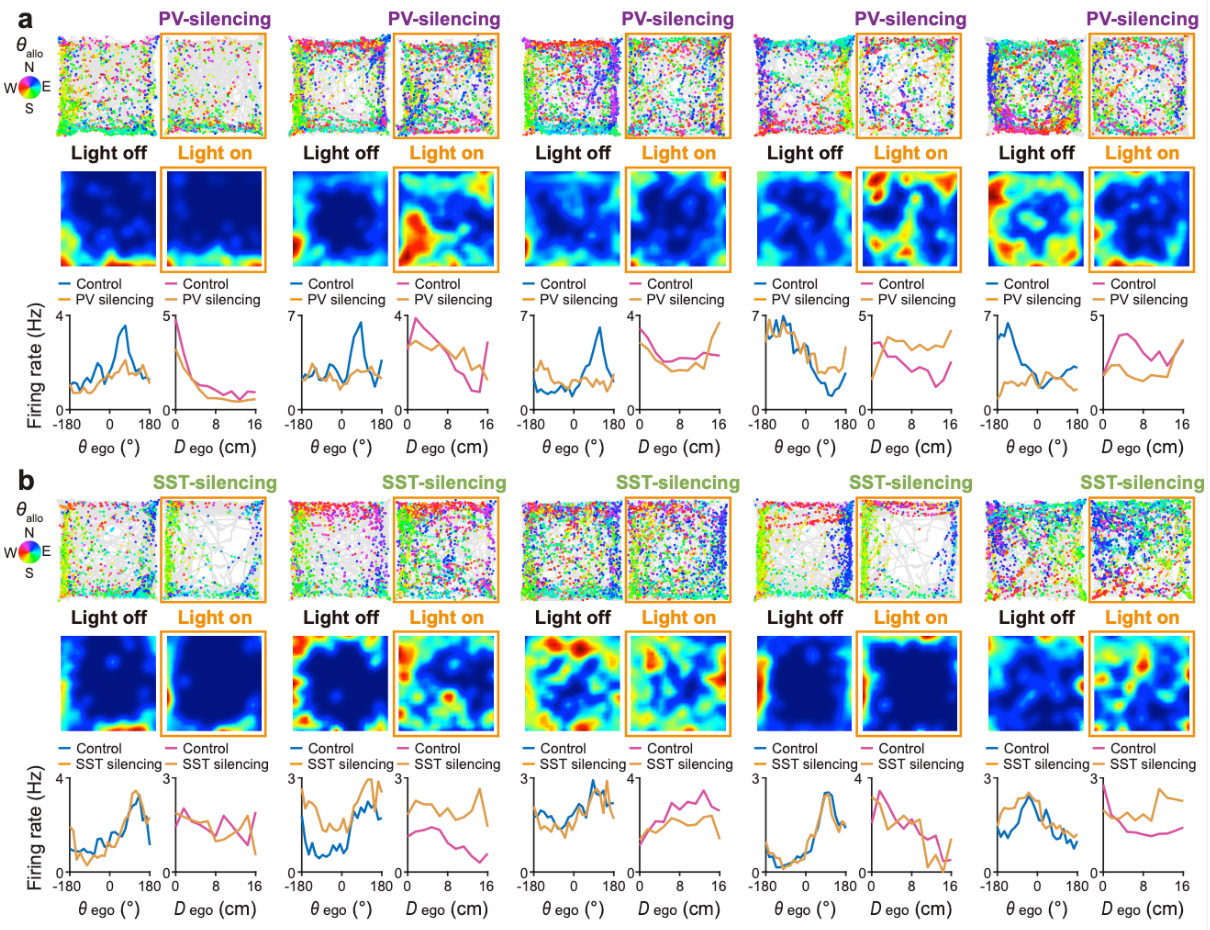


**Extended Data Fig. 5 Egocentric tuning of excitatory neurons during optogenetic silencing of PV and SST interneurons in retrosplenial cortex.**

**a,** Representative spike-trajectory plots (mouse trajectory: gray line, spike locations: colored dots) under the control (top left) and during optogenetic silencing of NpHR-expressing PV interneuron (top right, orange square border), with corresponding firing rate maps (middle, orange square border). Each spike is color-coded for allocentric head direction ($\theta_{allo}$) (inset, North: N; East: E; South: S; and West: W). (bottom left) Egocentric bearing ($\theta_{ego}$) tuning curves of RSC excitatory neurons under the control (dark blue) and during optogenetic silencing of NpHR-expressing PV interneurons (orange). (bottom right) Egocentric distance ($D_{ego}$) tuning curves under the control (magenta) and during optogenetic silencing of NpHR-expressing PV interneurons (orange).

**b,** Same as **a,** but for during optogenetic silencing of NpHR-expressing SST interneurons. n = 616 excitatory neurons from seven PV-Cre mice; n = 724 excitatory neurons from eight SST-Cre mice.

Extended Data Fig. 6

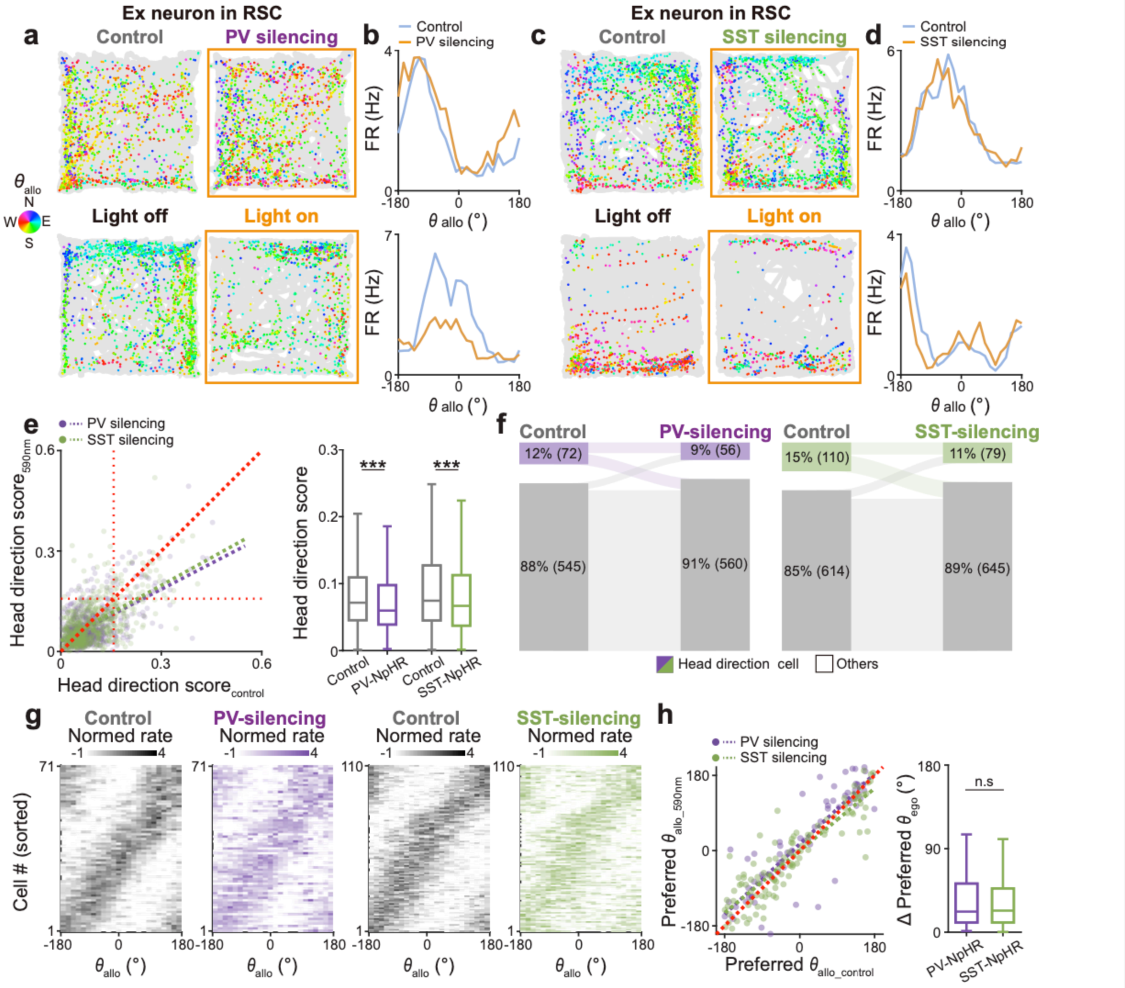


**Extended Data Fig. 6 Head direction tuning of excitatory neurons during optogenetic silencing of PV and SST interneurons in the retrosplenial cortex.**

**a,** Representative spike-trajectory plots (mouse trajectory: gray line, spike locations: colored dots) under the control (left) and during optogenetic silencing of NpHR-expressing PV interneurons (right). Each spike is color-coded for allocentric head direction ($\theta_{allo}$) (inset, North: N; East: E; South: S; and West: W).

**b,** Allocentric head direction ($\theta_{allo}$) tuning curves of RSC excitatory neurons in control (dark blue) and during optogenetic silencing of NpHR-expressing PV interneurons (orange).

**c-d,** Same as **a-b,** but with optogenetic silencing of SST interneurons.

**e,** (left) head direction score under optogenetic silencing of NpHR-expressing PV (purple) and SST interneurons (green) plotted as a function of head direction score under the control condition with linear regression fits (dotted line). (right) Box plots of head direction scores for each condition.

**f,** Sankey diagrams of head direction cell transitions during optogenetic silencing of NpHR-expressing PV (left) and SST interneurons (right).

**g,** Normalized $\theta_{allo}$ tuning maps of head direction cells under the control and during optogenetic silencing of NpHR-expressing PV and SST interneurons. Cells are ordered by preferred $\theta_{allo}$ in the control condition.

**h,** (left) Preferred $\theta_{allo}$ during optogenetic silencing of NpHR-expressing PV (purple) or SST interneurons (green) plotted as a function of preferred preferred $\theta_{allo}$ under control condition, with linear regression fits (dotted lines). (right)

Box plots showing the mean angular shift (Δ preferred $\theta_{allo}$) under optogenetic silencing of NpHR-expressing PV (purple) and SST interneurons (green). Box plot (**e**, **h**) shows 25th (lower box line), 50th (middle line), 75th (upper box line) percentile values and minimum and maximum values (whiskers). Paired t-test, $***P_{\text{Control-PV silencing}} = 2.70 \times 10^{-7}$ (**e**), $P_{\text{Control-SST silencing}} = 4.86 \times 10^{-6}$ (**e**). Two-sided Wilcoxon rank-sum test, $P = 0.6535$ (**h**).

n = 616 excitatory neurons from seven PV-Cre mice, n = 724 excitatory neurons from eight SST-Cre mice.

#### Extended Data Fig. 7

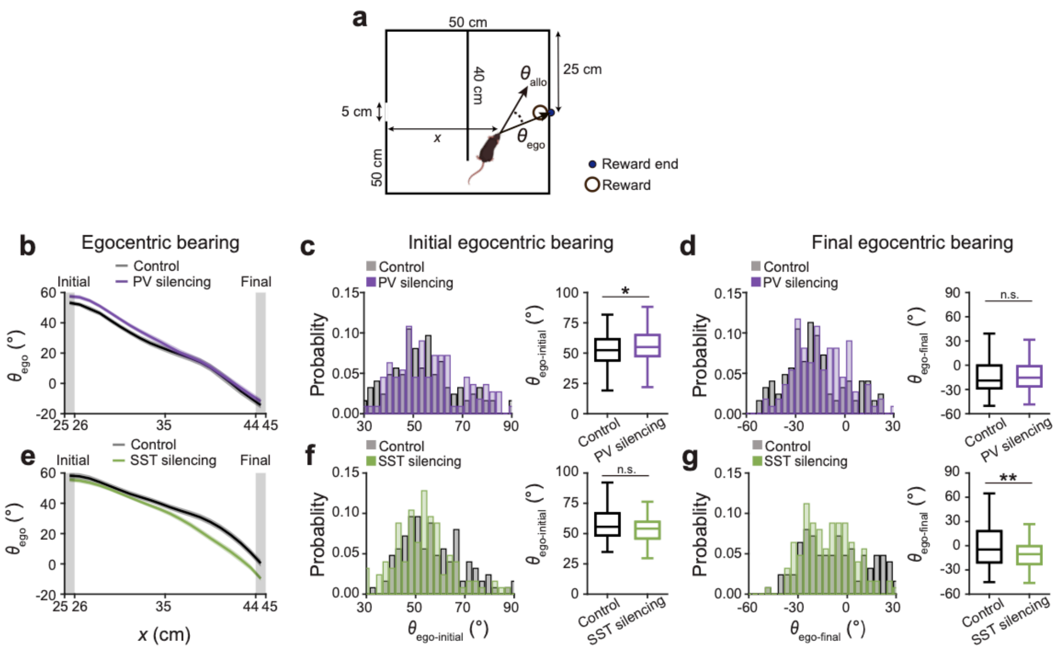


**Extended Data Fig. 7 The roles of PV and SST interneurons in guiding barrier-to-goal spatial navigation**

**a**, Schematic of the barrier-detour goal-directed navigation task showing the reward location (black circle), distance to start wall (*x*), the barrier gap end (blue dot), allocentric head direction ($\theta_{allo}$) and egocentric bearing to the reward end ($\theta_{ego}$) relative to the animal's instantaneous heading direction.

**b**, $\theta_{ego}$ toward the reward location as a function of *x* during the control condition (Day 4, black) and during PV interneuron silencing (purple). Solid lines indicate the mean and shaded areas represent mean ± SEM. Gray shaded regions indicate the first and the last bin.

**c,** (left) Probability distribution of $\theta_{ego\text{-}initial}$, defined as the mean $\theta_{ego}$ within the first spatial bin x = 25–26 cm, for control and PV interneuron silencing conditions. (right) Box plots of $\theta_{ego\text{-}initial}$ comparing control and PV interneuron silencing conditions.

**d**. Same as **c**, but for $\theta_{ego\text{-}final}$, defined as mean $\theta_{ego}$ when animals pass through x = 44–45 cm.

**e-g,** Same as **b-d,** but during optogenetic silencing of SST interneurons.

Box plots (**c**,**d**,**f**,**g**) show 25th (lower box line), 50th (middle line), and 75th (upper box line) percentile values; whiskers indicate minimum and maximum values. Watson-Wiliams test, $*P_{\text{Control-PV silencing}} = 0.0399$ (**c**), $P_{\text{Control-PV silencing}} = 0.1127$ (**d**), $P_{\text{Control-SST silencing}} = 0.3885$ (**f**), $**P_{\text{Control-SST silencing}} = 0.0080$ (**g**). n = 113 trajectories from five PV-Cre mice; n = 125 trajectories from five SST-Cre mice.
